# Extreme mobility creates contrasting patterns of panmixia, isolation-by-distance and hybridization in four flying-fox species (*Pteropus*)

**DOI:** 10.1101/2025.04.06.647431

**Authors:** Armin Scheben, Adam McKeown, Tom Walsh, David A. Westcott, Suzanne S. Metcalfe, Eric P. Vanderduys, Bruce L. Webber

## Abstract

Shifts between population connectivity and isolation shape evolutionary divergence, but their effects remain underexplored in highly mobile taxa. Flying-foxes (*Pteropus*) are a compelling model to study evolutionary divergence because of their rapid radiation in the island-rich region of the Indo-Australian Archipelago. Here, we investigate the population genetics of the highly mobile flying-foxes *Pteropus alecto*, *P. conspicillatus*, *P. poliocephalus*, and *P. scapulatus* across large parts of their overlapping ranges in Australia, Indonesia, and New Guinea. Using reduced-representation sequencing of 242 individuals, we examined the extent to which panmixia, isolation-by-distance, and hybridization are prevalent in these lineages. Phylogenetic, principal component and demographic analyses indicated isolation-by-distance in *P. alecto*, including an unexpected east-west genetic split between Australian populations, whereas *P. conspicillatus* showed weak differentiation between Australia and New Guinea and *P. poliocephalus* and *P. scapulatus* were panmictic across Australia. We further found evidence that infrequent hybridization and asymmetric gene flow, predominantly from *P. conspicillatus* to *P. alecto*, have shaped these recently diverged species. Although gene flow connected sampled populations along the geographic chain spanning Indonesia, Australia, and New Guinea, our results did not confirm the hypothesis that *P. alecto* and *P. conspicillatus* form a ring species. Rather, demographic analyses supported recent secondary contact during the last 14,848 to 35,856 years after a period of isolation between *P. alecto* and *P. conspicillatus*. Together, our findings indicate that long-range connectivity and spatially heterogeneous restrictions on gene flow have shaped the evolution and diversification of flying-foxes.

## 1 Introduction

Dispersal influences the evolutionary trajectory of species, both promoting speciation through geographic isolation and counteracting it by maintaining genetic connectivity (Lomolino, 2000). Through its effect on gene flow, range expansions, and colonization ability, dispersal plays a complex role in allopatric speciation, though speciation may also occur in the face of gene flow (Wu, 2001). Moreover, speciation can occur in sympatry due to non-geographic barriers such as ecological divergence (Richards et al., 2019). Because dispersal is an important driver of gene flow across space, many species capable of long-distance dispersal form panmictic populations across continents (Bruxaux et al., 2024; Neethling et al., 2008; Peel et al., 2013; Reich et al., 2024; Reudink et al., 2011). In contrast, genetic isolation-by-distance is also widespread, leading to correlations between populations’ geographic structure and their genetic structure (Aguillon et al., 2017; Sharbel et al., 2000; Wright, 1943). Panmixia and isolation-by-distance can occur within species at various spatiotemporal scales and incipient speciation can be prevented by secondary contact if populations are not sufficiently reproductively isolated (Karin et al., 2025; Seehausen et al., 2008; Vonlanthen et al., 2012). Alternatively, secondary contact can lead to reinforcement of species boundaries through hybrid inviability, and long-lasting natural hybrid zones can even be maintained despite gene flow across large geographic distances (Abbott et al., 2013). In rare cases, reproductively isolated populations connected by interbreeding populations around geographic barriers may form “ring species” (Irwin et al., 2005). Although ideal ring species may not exist, ‘broken’ ring species without continuous gene flow during divergence can still provide insights into ringlike evolutionary dynamics (Alcaide et al., 2014; Kuchta & Wake, 2016). However, our understanding of these complex stages of dispersal and speciation remains limited by how infrequently these processes have been investigated in recently diverged, highly mobile species.

Bats are the only mammals capable of powered flight and make up the second-most species-rich mammal group with 1,487 species (Simmons & Cirranello, 2025). Flying-foxes of the genus *Pteropus* (Pteropodidae: Chiroptera) are a particularly fast-radiating and recently diverged genus of bats originating in the Wallacea region of the Indo-Australian Archipelago (Tsang et al., 2019). The rapid diversification of flying-foxes has been associated with their high long-distance dispersal rate compared to other volant species (Tsang et al., 2019), enabling them to colonize new and isolated habitats such as archipelagos, where subsequent geographic isolation promotes allopatric speciation (Smyčka et al., 2023). Individual flying-foxes are capable of long-distance movements (Breed et al., 2010; Eby, 1991; Westcott et al., 2015), with tracking experiments showing some individuals covering over 100 km a day (Roberts et al., 2012) and an average of 1,427–6,073 km annually (Welbergen et al., 2020). Flying-fox biogeography reflects this extreme mobility, with repeated colonizations of remote oceanic islands (O’Brien et al., 2009). While all Australian flying-fox species are known to make long-distance movements, the extent and frequency of connectivity between different parts of species’ ranges, how this is influenced by geography, and the evolutionary consequences are virtually unknown.

Four flying-fox species are native to the Australian mainland: the black flying-fox (*Pteropus alecto* Temminck, 1837), spectacled flying-fox (*P. conspicillatus* Gould, 1850), grey-headed flying-fox (*P. poliocephalus* Temminck, 1825), and little-red flying-fox (*P. scapulatus* Peters, 1862). While *P. poliocephalus* is endemic to Australia, the other species have disjunct distributions extending to Indonesia and New Guinea (Figure 1). Four subspecies of *P*. *alecto* have been described including *P*. *alecto gouldi* from Australia and New Guinea, *P. alecto aterrimus* from the Bawean and Kangean islands, *P*. *alecto morio* from Sawu and Sumba, and *P*. *alecto alecto* from Lombok, Salayar, and Sulawesi (Mickleburgh et al., 1992). These subspecies are poorly studied and may not be monophyletic (Tsang et al., 2019). An early genetic study found that the widespread Australian *P. alecto*, *P. poliocephalus*, and *P. scapulatus* were panmictic, suggesting these bats resembled migratory species (Webb & Tidemann, 1996). The less widely distributed *P. conspicillatus* has a notably smaller population size compared to the other species and is listed as endangered in Australia (Roberts et al., 2020). Previous work has hypothesized that hybridization occurs between Australian *P. alecto* and both *P. conspicillatus* (Fox, 2006; Neaves et al., 2018) and *P. poliocephalus* (Webb & Tidemann, 1995) based on limited genetic evidence. *P. alecto* and *P. conspicillatus* are morphologically highly similar, with the key distinguishing morphological features of *P. conspicillatus* being its dentition and eye-rings, though light eye-rings also occur in *P. alecto* (Neaves et al., 2018). Past phylogenetic studies of the relationship between *P. alecto* and *P. conspicillatus* using small sets of nuclear and mitochondrial markers have often found mitonuclear discordance and paraphyly (Almeida et al., 2014; Fox, 2006; Tsang et al., 2019), explaining this through incomplete lineage sorting or hybridization (Almeida et al., 2014). The divergence-time of these two species is variously estimated at 1–2 million years ago (mya) (Tsang, 2015) or 2–3 mya (Tsang et al., 2019). Intriguingly, their geographic distribution across Australia, Indonesia, and New Guinea suggests that *P. alecto* and *P. conspicillatus* may represent a ring species.

**Figure 1.**
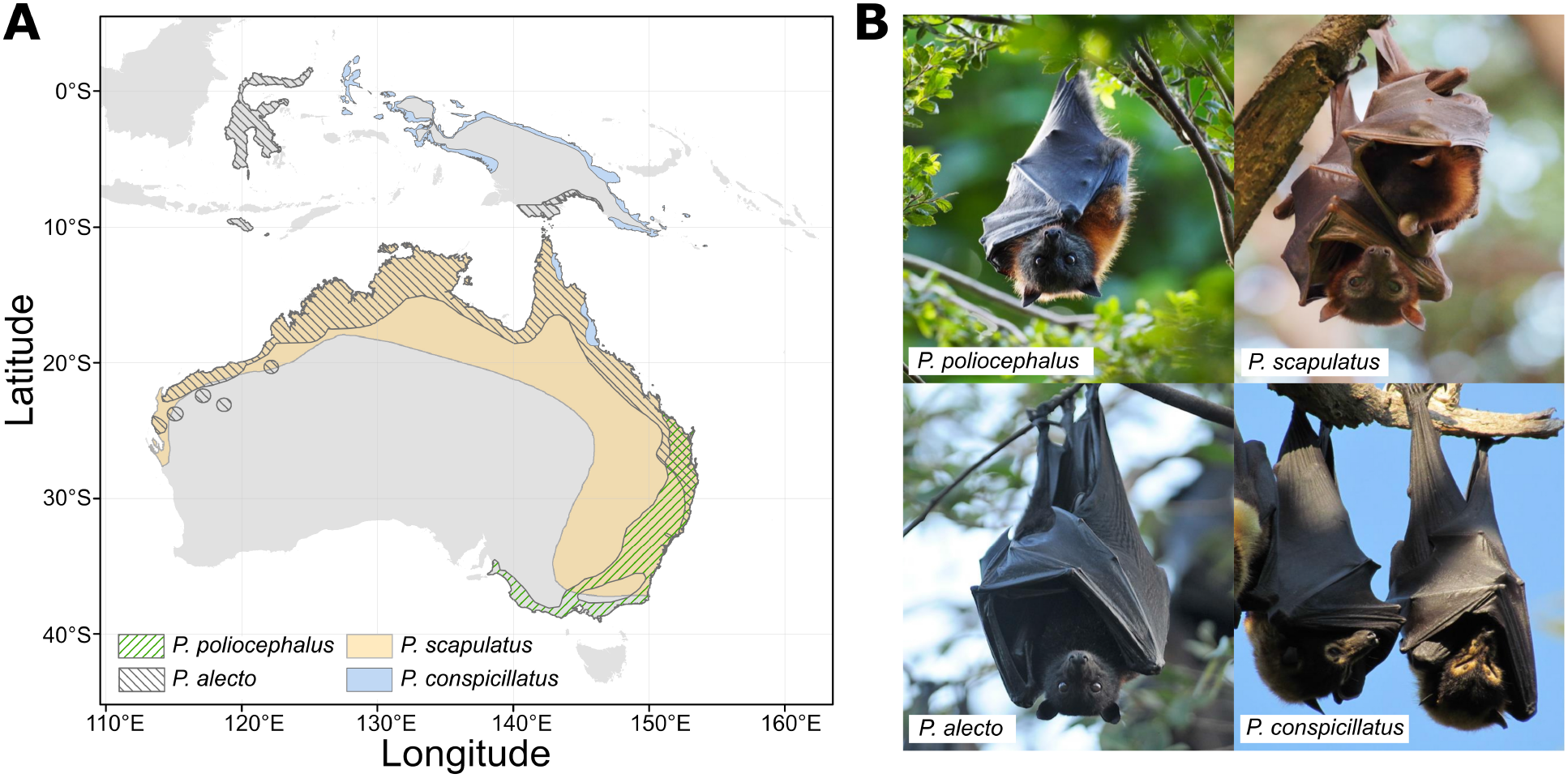
Sampled *Pteropus* flying-fox species and their geographic distributions. A) Ranges of the four flying-fox species native to the Australian mainland (https://www.iucnredlist.org). B) Photographs of the four flying-fox species. Image credit: Adam McKeown and Eric P. Vanderduys.

In this work, we investigate the population genetic structure of *P. alecto*, *P. conspicillatus*, *P. poliocephalus*, and *P. scapulatus* using genome-wide SNPs. We aim to determine 1) how the genetic structure of these four flying-fox species differs across their distributions, 2) what role hybridization has played in their evolution, and 3) whether *P. alecto* and *P. conspicillatus* form a ring species. We hypothesize that mainland Australian flying-fox populations are panmictic, but that Indonesian populations of *P. alecto* and New Guinean populations of *P. conspicillatus* are genetically distinct from Australian populations. In addition, we hypothesize that *P. alecto* hybridizes with the related flying-fox species *P. conspicillatus* and *P. poliocephalus* in regions of overlapping distribution. Finally, we hypothesize that *P. alecto* and *P. conspicillatus* form an interbreeding population with gradual isolation-by-distance along the geographic ring connecting Australia with Indonesia, and New Guinea, with the endpoint in Central Indonesia.

## 2 Materials and methods

### 2.1 Sample collection

A total of 288 samples from individuals of the four different flying-fox species, *P. alecto* (n = 125), *P. poliocephalus* (n = 47), *P. scapulatus* (n = 51), and *P. conspicillatus* (n = 65) were collected from 27 roost locations in Australia and New Guinea as well as 4 locations in Indonesia (Table S1). The animals were captured in accordance with animal ethics approval (Permit number CSIRO AEC 2016-17). Small 3 mm circles of skin were removed from the wing membrane of individual flying-foxes using a biopsy punch. The samples also included 33 historical *P. alecto* specimens collected between 1989 and 2003 from the Western Australian museum collection (WAM). Preservation conditions for historical samples were not available. We analyzed the historical and extant specimens together because we do not expect the age differences of 10–29 years (2–7 generations) between these specimens to substantially impact our inferences at evolutionary time scales.

### 2.2 Genomic library preparation, sequencing, and variant calling

Genomic DNA was extracted using the Qiagen DNA Blood and Tissue Kit (Hilden, Germany). A total amount of 100 ng of genomic DNA from each individual was digested in a 50 µL reaction with 20 units each of high-fidelity restriction enzymes, PstI and EcoRI (New England Biolabs, Ipswich MA, USA) and a sequencing library prepared (see Text S1 for additional details). Sequencing was conducted using the 75 bp PE v3 sequencing kit on an Illumina NextSeq. Read quality control was conducted using FastQC (http://www.bioinformatics.babraham.ac.uk/projects/fastqc/). Demultiplexing and trimming to 43 bp was carried out using stacks 2.52 process_radtags (Rochette et al., 2019). DNA degradation was assessed with mapDamage 2.2.2 (Jónsson et al., 2013). Reads were aligned to the *P. alecto* reference genome ASM32557v1 (Zhang et al., 2013) from GenBank using BWA-MEM 0.7.17-r1198-dirty (Li, 2013) with default settings. The low-quality 5 bp flanking regions of all reads were clipped using GATK 4.6.2.0 (Van der Auwera & O’Connor, 2020). Using BCFtools 1.19 (Danecek et al., 2021), a pileup was generated with a minimum base quality of 10 and minimum mapping quality of 20 and variants were called, applying Hardy-Weinberg equilibrium assumptions within populations using the -G flag. Invariant sites were included to inform demographic modeling. Following variant calling, pre-analysis quality control of alignment metrics, preliminary phylogenetic species assignment, and duplicate removal were conducted, excluding 46 samples (Text S2 and Table S1). Based on this analysis, for all analyses below we split *P*. *alecto* into *P*. *alecto alecto*, which includes phylogenetically deeply diverged samples primarily from Sulawesi, and the remaining *P*. *alecto*. Variant calling using BCFtools with the same parameters used initially was then repeated using the final set of 242 samples comprising *P. alecto* (n = 86), *P. alecto alecto* (n = 9), *P. poliocephalus* (n = 45), *P. scapulatus* (n = 47), and *P. conspicillatus* (n = 55).

The variant calls were split into three datasets for filtering and analysis (Table S2): an “all” set containing all four species (n = 242), an “alecto + conspicillatus” set (n = 150), and an “alecto ex alecto alecto + conspicillatus” set (n = 141) which excludes *P*. *alecto alecto*. Filtering was conducted separately for these datasets to minimize the number of SNPs excluded due to missingness in species-specific analyses. Sites were filtered using VCFtools 0.1.16 (Danecek et al., 2011) using a minimum depth of 5, a minimum genotype quality of 20, a minimum minor allele frequency (MAF) of 0.05, and a maximum missingness of 20% across all samples. Next, variants were filtered to retain only biallelic SNPs. PLINK2 2.00a2.3LM (Chang et al., 2015) was used to prune linked SNPs with “--indep-pairwise 50 5 0.3”. Kinship coefficients were calculated using the KING algorithm (Manichaikul et al., 2010) as implemented in PLINK2 and duplicates as well as first-degree relatives were excluded based on a filter of 0.2 because first-degree relatives are expected to have a coefficient of 0.25.

Modified SNP filtering parameters were used for two analyses. Firstly, for genome-wide heterozygosity and nucleotide diversity estimates, we only applied the above thresholds for minimum depth, minimum genotype quality, and maximum missing genotypes, retaining sites that have low MAF, are multiallelic, or are in linkage disequilibrium to ensure unbiased counts of variable sites and rare alleles. Secondly, for fastsimcoal2 modeling, we used SNPs filtered using the above thresholds for minimum depth, minimum genotype quality, maximum missing genotypes and biallelic sites, but we did not filter for linkage disequilibrium or MAF. We retained linked SNPs to avoid biasing our calculated proportion of polymorphic sites and to maximize information for fastsimcoal2 parameter estimation, which is robust to the inclusion of these SNPs in the site frequency spectrum. Similarly, we did not filter rare alleles by MAF because they are informative for site frequency spectrum analysis.

### 2.3 Population genetic analysis

By investigating genetic diversity, relatedness, and admixture, we aimed to characterize the genetic differentiation, connectivity, and potential hybridization among *Pteropus* species across their geographic ranges. For this population genetic investigation, individuals were assigned to six geographic regions: Indonesia, North West Australia, North East Australia, East Australia, Wet Tropics Australia, and New Guinea (Figure S1). The Wet Tropics region of Queensland, Australia is a unique region of wet tropical forests and distinct from other sampled locations in North East Australia. We then assessed nucleotide diversity in non-overlapping 100 kb windows using pixy 1.2.11.beta1 (Korunes & Samuk, 2021), removing windows with fewer than 50 sites. We calculated the genome-wide heterozygosity (*Het*) per individual using BCFtools as the proportion of callable sites that were heterozygous. Diversity was compared between species and populations using pairwise Wilcoxon rank-sum tests.

Using both the “all” and the “alecto + conspicillatus” SNP sets, we calculated genome-wide genetic differentiation (F_ST_) for population pairs using stacks populations, carried out principal component analyses (PCA) with SNPRelate 1.40.0 (Zheng et al., 2012), inferred phylogenetic trees with RAxML 8.2.12 (Stamatakis, 2014) using the GTRCAT model and 100 bootstrap replicates, and, because the assumption that all sites share a genealogical history may not hold, reconstructed phylogenies using the coalescent approach CASTER-site 1.23.2.6 (Zhang et al., 2025). We also used both SNP sets to investigate between-species divergence times with SNAPP (Text S3). To determine whether F_ST_ was significantly different from 0, we permutated population labels and recalculated F_ST_ for 1,000 permutations.

For all further analyses we focused on *P*. *alecto* and *P*. *conspicillatus*, relying on the “alecto + conspicillatus” SNP set or, where *P. alecto alecto* was not included, the “alecto ex alecto alecto + conspicillatus” SNP set. To detect admixture and potential hybridization, population structure was inferred using fastSTRUCTURE 1.0 (Raj et al., 2014) with 1 to 15 clusters and the default prior, using the provided chooseK method to select an optimal number of clusters that maximizes the log-marginal likelihood lower bound of the data. Admixture proportions for individuals were estimated as continuous values representing membership fractions across clusters, with no fixed threshold for cluster assignment, and clusters were interpreted qualitatively by linking them to species and geographic regions. To validate potential hybridization inferred using admixture analysis, we selected ancestry-informative markers with a species-specific allele frequency of at least 0.85 and used them to generate triangle plots of hybrid index and interclass heterozygosity using triangulaR (Wiens & Colella, 2024). Individuals with over one third of ancestry-informative markers missing were excluded from the analysis.

### 2.4 Gene flow inference

To elucidate the extent and direction of historical and contemporary gene flow between genetic populations of *Pteropus alecto* and *P. conspicillatus*, we used a set of three complementary analyses. Firstly, migrations between the genetic populations of *P*. *alecto* and *P*. *conspicillatus* were inferred using TreeMix 1.13 (Pickrell & Pritchard, 2012) with the Indonesian *P. alecto alecto* individuals set as the outgroup. We used *P. alecto alecto* as the outgroup because these samples represent the most diverged taxon in our dataset. TreeMix uses SNP-based allele frequency data to infer discrete migration events, which are modelled as migration edges *m* between populations on a phylogenetic tree. We ran TreeMix with 500 SNPs per block for covariance matrix estimation and values of *m* ranging from 0 to 7, with 10 replicates for each value. We then used the Evanno method implemented in the R package OptM to determine the optimal value of *m* (Fitak, 2021). We next computed an additional 100 replicates using the optimal value of *m* and selected the replicate with the highest likelihood.

Secondly, to complement the TreeMix analysis by also explicitly exploring spatial genetic structure, we applied FEEMSmix 2.0.0 to calculate an effective migration surface (Shastry et al., 2025). FEEMSmix adds long-range migration to the FEEMS framework, which infers migration rates on the edges of a graph of connected nodes representing local populations. Cross-validation was used to identify the tuning parameters comprising the migration edge weight penalty (*λ* = 0.05) and the node-specific variance penalty (*λ_q_* = 100). These tuning parameters penalize the strength of migration by homogenizing neighboring nodes and edges.

Finally, to better model continuous gene flow across different time periods, we investigated gene flow between and within the *P*. *alecto* and *P*. *conspicillatus* populations using coalescent simulations based on the site frequency spectrum implemented in fastsimcoal2 2.8 (Excoffier et al., 2013). To account for missing data, we maximized the number of segregating sites by using a projection of samples based on EasySFS (https://github.com/isaacovercast/easySFS) which relies on methods implemented in dadi (Gutenkunst et al., 2009). The “--total-length” parameter was set based on the variant and the invariant sites detected in the aligned reads for each population pair. We defined four demographic scenarios for comparison between population pairs: no migration, ancient migration, recent migration, and constant migration (Figure 2). We allowed for asymmetric migration between populations. We assumed a mutation rate of 2.5×10^-8^ mutations per nucleotide site per generation based on robust calculation in humans (Nachman & Crowell, 2000; Sovic et al., 2016) and a generation time of 4 years (McIlwee & Martin, 2014). To test the impact of our assumed human mutation rate, we also tested an average mammalian rate of 2.2×10^-9^ per nucleotide site per year (Kumar and Subramanian, 2002) multiplying it with the *Pteropus* generation time of 4 years to obtain an alternative mutation rate of 8.8×10^-9^ per nucleotide site per generation. To evaluate the demographic scenarios, we ran fastsimcoal2 for 100 replicates per migration scenario using the parameters “-m -n 50000 -L 30 -M -C 10”. After selecting the replicate with the highest maximum composite likelihood for each scenario, we used the Akaike information criterion (AIC) to determine the best fitting migration scenario for each population pair, assessing consistency with the corrected AIC (AICc). Next, to calculate confidence intervals for all model parameter estimates, we ran 100 parametric bootstrap replicates by simulating site frequency spectra from the estimates of each of the best-fitting models (Excoffier et al., 2013).

**Figure 2.**
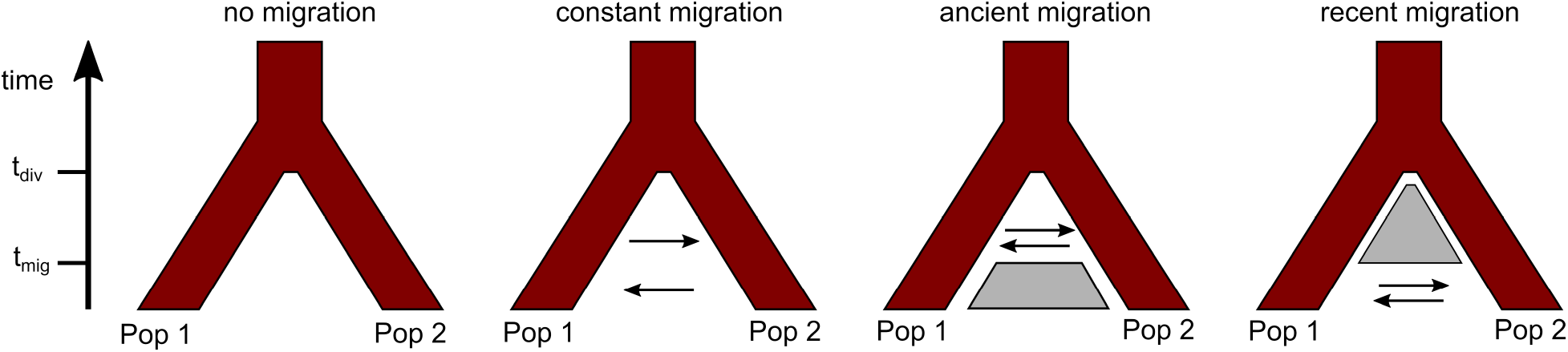
Migration models for pairwise demographic histories between populations. Each model includes a parameter for the time that populations split (t_div_) and models with non-constant migration include a parameter for the time that migration began or ceased (t_mig_) depending on whether the migration occurred recently or in the past. Grey boxes indicate barriers to gene flow.

## 3 Results

### 3.1 Sequencing, read processing, quality control, and variant calling

A total of 242 of 288 samples were retained following quality control (Table S1). The passing samples did not include any first-degree relatives. For each sample passing quality control, an average of 2.14 million read pairs (SD = 0.79 million) of 43 bp trimmed length were generated. The average alignment rate to the *P. alecto* reference genome was 95.12% (SD = 1.45%), covering an average of 4.39% (SD = 1.62%) of the genome and 7.95% (SD = 3.11%) of the coding regions with an average 1.8-fold (SD = 0.13) overrepresentation of coding regions (Table S1). The average per species percentage of covered regions that were coding took on a narrow range of 6.56–7.51%. Alignment rates did not differ between *P*. *alecto* and *P*. *conspicillatus* (mean = 96.2%, unpaired *t*-test Holm-adjusted *p* = 1) but alignment rates for the more diverged species *P*. *poliocephalus* (mean = 93.1%, unpaired *t*-test Holm-adjusted *p* = 1.59×10^-50^) and *P*. *scapulatus* (mean = 94.3%, unpaired *t*-test Holm-adjusted *p* = 6.26×10^-24^) were slightly lower. A total of 1,339,950 raw variants were detected and then filtered for minimum allele depth, missingness per variant, minor allele frequency, and linkage disequilibrium (see Methods). After filtering, the “all”, “alecto + conspicillatus”, and “alecto ex alecto alecto + conspicillatus” datasets retained 11,818, 17,708, and 18,285 SNPs with average per-sample depths of 17.90×, 16.83×, and 16.96×, respectively (Table S2). The samples from the WA museum showed a mean genotype missingness rate of 13.4% which did not differ from the extant samples (unpaired *t*-test *p* = 0.20). DNA degradation based on cytosine deamination, as measured by C to T substitution rates, also did not differ between museum and extant samples (unpaired *t*-test *p* = 0.45; Figure S2).

### 3.2 Population structure of four flying-fox species

We found a wide range of genome-wide diversity in the flying-fox species as measured by individual *Het* (Figure S3, Tables S3 and S4) and nucleotide diversity *π* (Figure S4, Tables S5 and S6). In between-species comparisons, each species exhibited significant differences in *Het* (Holm-adjusted *p* ≤ 1.84×10^-5^) and *π* (Holm-adjusted *p* ≤ 4.40×10^-19^) based on two-sided pairwise Wilcoxon rank-sum tests. We found intermediate diversity relative to the other species in *P. alecto* (mean *Het* = 3.77×10^-3^, mean *π* = 3.51×10^-3^) and *P. conspicillatus* (mean *Het* = 3.42×10^-3^, mean *π* = 3.14×10^-3^). *P*. *scapulatus* had the highest diversity (mean *Het* = 5.65×10^-3^, mean *π* = 5.26×10^-3^) and the lowest diversity values were found in *P*. *poliocephalus* (mean *Het* = 3.16×10^-3^, mean *π* = 2.84×10^-3^) and *P*. *alecto alecto* (mean *Het* = 2.52×10^-3^, mean *π* = 2.24×10^-3^). Within *P. alecto*, Australian populations showed no differences in *Het* (Holm-adjusted *p* ≥ 8.60×10^-1^) but showed elevated *Het* compared to Indonesian population (Holm-adjusted *p* ≤ 5.36×10^-4^) based on two-sided pairwise Wilcoxon rank-sum tests. Similarly, nucleotide diversity was significantly higher in Australian *P*. *alecto* compared to Indonesian *P*. *alecto* (two-sided pairwise Wilcoxon rank-sum test, Holm-adjusted *p* ≤ 5.58×10^-158^).

Across the four flying-fox species, we observed a wide range of between-species genetic differentiation, with F_ST_ ranging from 0.06 for *P*. *alecto*–*P*. *conspicillatus* to 0.41 for *P*. *poliocephalus*–*P. alecto alecto* (Figure S5, Table S7). Genetic differentiation between all compared species and populations was significantly greater than 0 (permutation tests with 1,000 replicates, *p* ≤ 0.001). Within the closely related *P*. *alecto*–*P*. *conspicillatus* pair (Figure S6, Table S8), F_ST_ values indicated low genetic differentiation between the New Guinean and the Australian *P*. *conspicillatus* (F_ST_ = 0.01) but higher differentiation between the Indonesian and Australian *P*. *alecto* populations (F_ST_ = 0.05-0.06). Additionally, we found that the Indonesian *P*. *alecto* population was most differentiated from the New Guinean *P*. *conspicillatus* (F_ST_ = 0.14) and less differentiated from the Australian *P*. *conspicillatus* (F_ST_ = 0.11). These trends were consistent with genetic differentiation calculated for genetic populations (Tables S9 and S10).

PCA delineated the samples by species, but suggested close genetic association between *P*. *alecto* and *P*. *conspicillatus* (Figure 3, Table S11). Although *P*. *conspicillatus* showed limited population structure, *P*. *alecto* individuals formed mostly discrete groups by country as well as showing some separation within Australia between North West Australia and the remaining groups in East Australia and North East Australia (Figure 3D, Table S12). In *P*. *alecto*, the first and fourth principal components reveal that all Burketown samples cluster with the North West Australia group and three North West Australian samples cluster with the Indonesia group (Figure S7). Based on population structure inference, the majority of Australian *P*. *alecto* individuals also show signs of admixture, either from the Indonesian *P*. *alecto* cluster or from the *P*. *conspicillatus* cluster (Figure 3E). *P. alecto* individuals such as sample 1702005 and also *P. conspicillatus* individuals such as sample 1306014 show substantial admixture (Table S13). As expected, these hybrid samples also show among the highest genome-wide heterozygosity. Sample 1306014 (*Het* = 4.16×10^-3^) has the second highest genome-wide heterozygosity in *P. conspicillatus* and sample 1702005 (*H*et = 4.33×10^-3^) has the second highest genome-wide heterozygosity in *P. alecto*. With an admixture proportion of 47.7% from the *P. alecto* cluster, the *P. conspicillatus* sample 1306014 is likely a hybrid or backcross.

**Figure 3.**
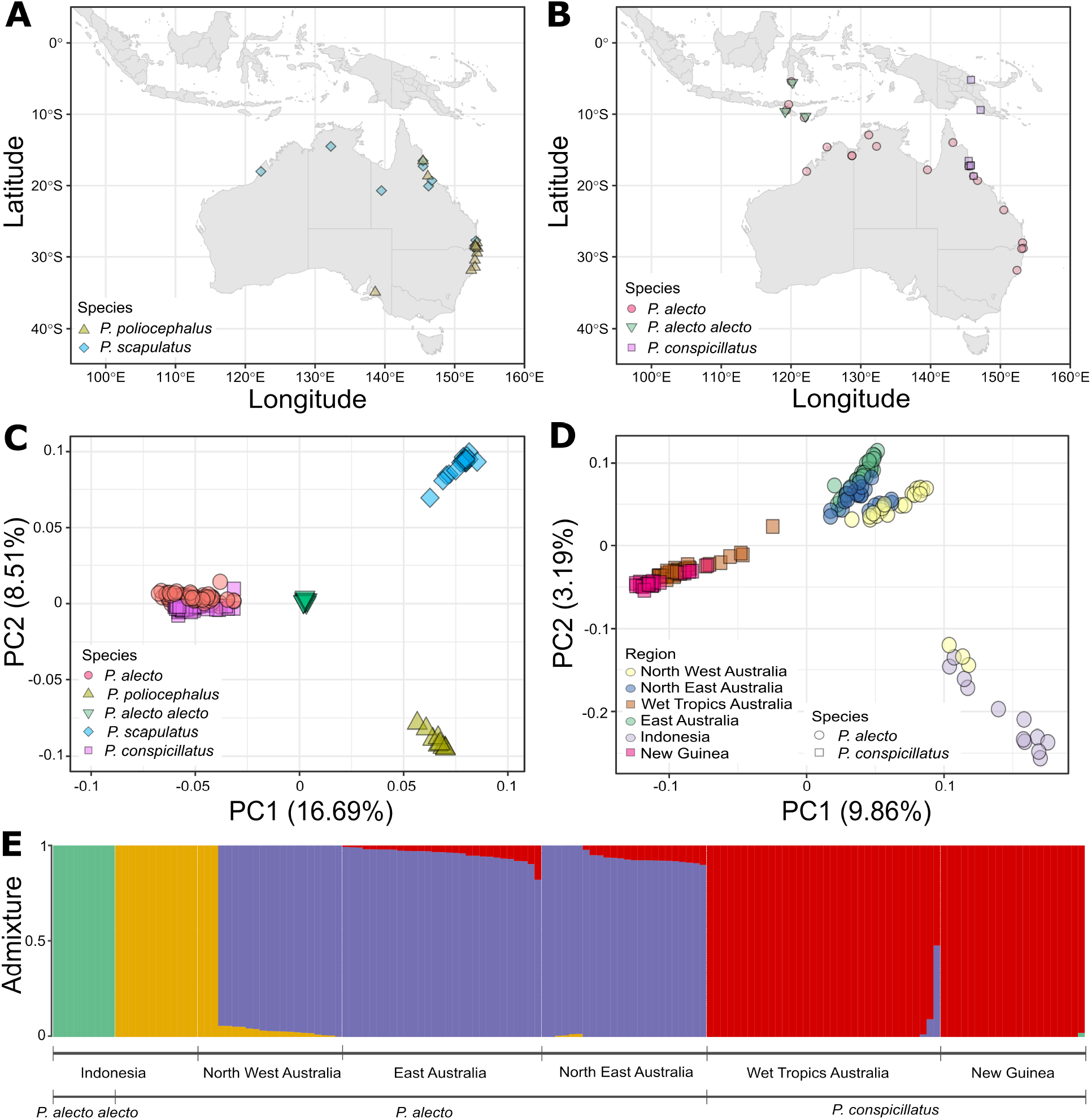
Population structure of four flying-fox species in Australia, Indonesia, and New Guinea. A) Sampling locations for *Pteropus poliocephalus* and *P*. *scapulatus*. B) Sampling locations for *P*. *alecto* and *P*. *conspicillatus*. C) SNP-based principal component analysis (PCA) of all sampled species. The percentage of genetic variance explained by each PC is shown on the axis. D) SNP-based PCA of six geographic populations of *P*. *alecto* and *P*. *conspicillatus*. The individual intermediate between the *P*. *conspicillatus* and the main Australian *P. alecto* cluster is sample 1306014. E) Structure plot of *P*. *alecto* and *P*. *conspicillatus* samples with four admixture groups. Bars indicate the proportion of each sample assigned to each group, revealing admixture between groups.

Missingness can spuriously pull samples to the center in PCA (Yi & Latch, 2022), but the missingness of 10.0% in sample 1306014 is average for the population (mean = 10.0%, SD = 9.5%). Furthermore, we did not find an effect of more relaxed genotype missingness thresholds on the PCA results (Figure S8). We also confirmed that the clustering of Indonesian and North West Australian samples from the WA museum was not related to biases in genotype missingness by conducting an additional PCA excluding all SNPs with missing genotypes (Figure S9). An evaluation of the effect of genotype missingness thresholds on the admixture analysis also found high consistency (Pearson’s correlation > 0.99) in per-sample admixture proportions between a filter of up to 20% missingness and filters of up to 50%, 40%, and 30% (Table S13).

PCA and model-based population structure inference cannot readily distinguish admixture from isolation-by-distance and may thus not be sufficient to infer hybridization (Wiens & Colella, 2025). We therefore calculated hybrid indices of all *P. alecto* and *P. conspicillatus* individuals, representing the proportion of ancestry received from each parental population based on a set of 15 ancestry-informative markers. The triangle plot based on hybrid index and interclass heterozygosity confirmed hybridization between the species, indicating that individual 1306014 is a *P. alecto* × F_1_ cross (Figures S10 and S11). In addition, *P. alecto* individuals including 1610001, 1702007, and 1702026 as well as *P. conspicillatus* individual 37721 were identified as later backcross generations. Overall, signatures of admixture from the *P*. *conspicillatus* cluster to the *P*. *alecto* cluster are more prevalent, with the majority of *P*. *conspicillatus* individuals not showing evidence of admixture or hybridization.

A SNP-based maximum likelihood phylogeny of *P*. *alecto* and *P*. *conspicillatus* recovered robust (bootstrap support > 70) clades grouping these two species as distinct from the outgroup, but none of the six geographic groups except Indonesia formed robust clades (Figure S12). The Indonesian *P*. *alecto* samples form a robustly supported clade with four North West Australian samples including three historical samples (M54660, M53754, and M54659) also clustered with Indonesian samples in the PCA and an extant individual (1702031). In contrast to *P*. *alecto* and *P*. *conspicillatus*, *P*. *poliocephalus* and *P*. *scapulatus* showed no geographic structure across the sampled populations (Figure S13). Although the genealogical history of the samples may violate some of the assumptions of the maximum likelihood phylogeny, the coalescent phylogenies were consistent with the maximum likelihood phylogenies (Figures S14 and S15).

### 3.3 Admixture occurs between *P. alecto* and *P. conspicillatus* populations along a geographic ring

We next investigated population splits and admixture events in more depth using a maximum likelihood approach to infer migration edges in a population tree and coalescent simulations based on the site frequency spectrum to infer migration rates. We grouped geographic populations based on the inferred genetic structure to avoid unsampled populations impacting our demographic estimates (Slatkin, 2005). Based on our PCA, we treated *P. conspicillatus* as a single population and *P. alecto* as three populations comprising 1) Indonesian samples and the three North West Australian samples clustered with them, 2) the remaining North West Australian and Burketown samples, and 3) the East Australian and remaining North East Australian samples (Figure S7). Although F_ST_ between North West Australian and the other Australian samples was significant, our population structure analyses did not conclusively support a discrete North West Australian population. Besides our primary grouping, we therefore also tested grouping all Australian samples except those clustered with Indonesian samples. Finally, to determine which role these three North West Australian samples clustering with Indonesian samples played in our results, we performed analyses excluding them. Unless otherwise stated, the results below refer to the primary grouping with three *P. alecto* populations. With these groupings, we then sought to show whether migration links the chain of populations between Indonesia, North West Australia, North East Australia and East Australia, and New Guinea (Figure S1).

We found support for two migration edges connecting *P. conspicillatus* to both Australian *P. alecto* populations (Figure S16A). When grouping Australian *P*. *alecto* genetic groups together, we found a single migration edge connecting *P. conspicillatus* to Australian *P*. *alecto* (not shown). At a geographic level, mean admixture proportions in roost locations are consistent with historical connectivity over substantial distances from Indonesia to North West Australia and from New Guinea and the Wet Tropics to East Australia (Figure 4A). The inferred effective migration surface emphasized high levels of within-species connectivity, including one weakly supported long-range migration from North East Australia to North West Australia and one from Indonesia to Australia (Figure 4B).

**Figure 4.**
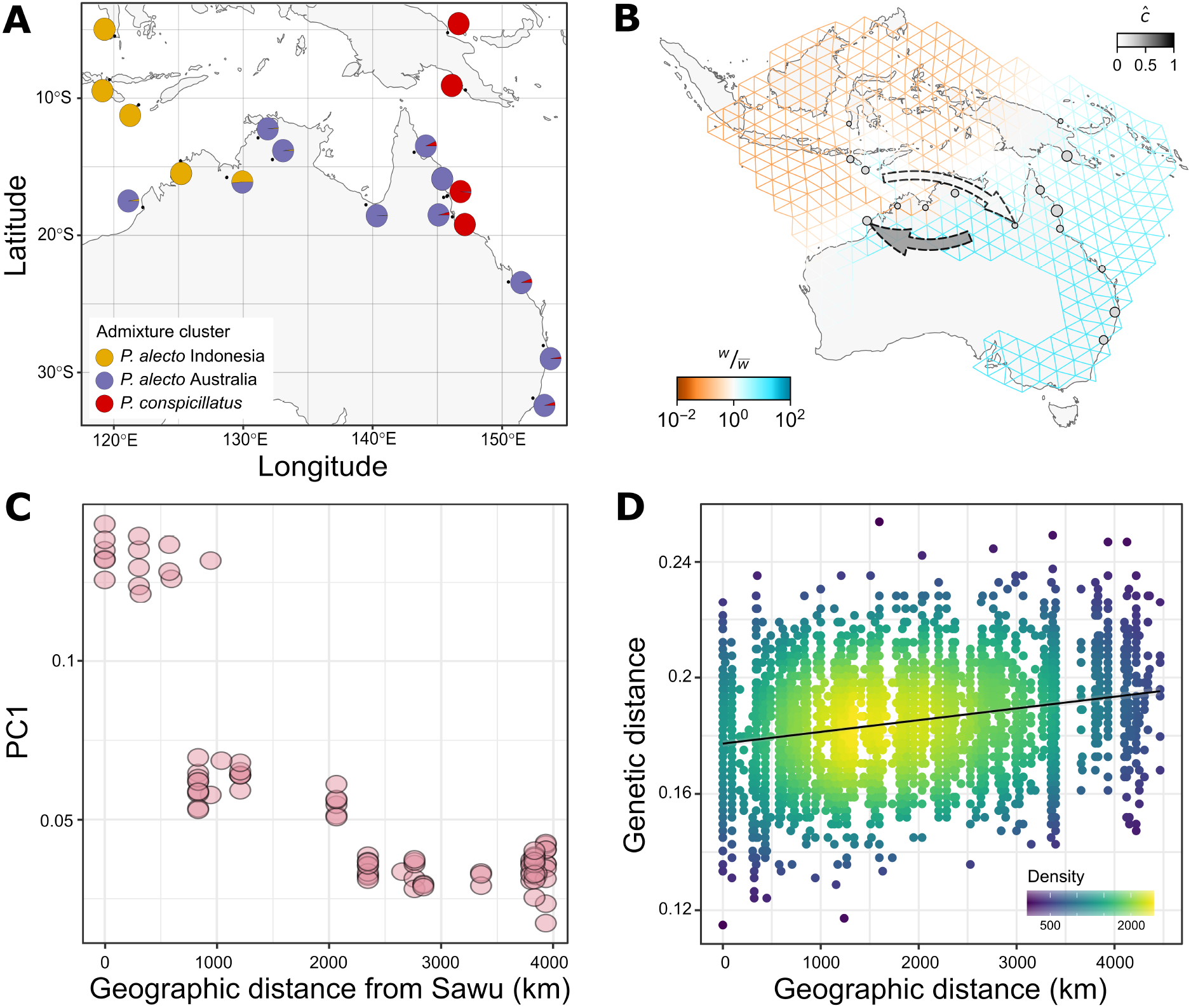
Gene flow in *Pteropus alecto* and *P. conspicillatus* along a geographic ring ranging across Indonesia, Australia, and New Guinea. A) Mean admixture proportions for local demes of *P. alecto* and *P. conspicillatus*. Admixture is inferred via fastSTRUCTURE (see Figure 3E). B) Effective migration surface with long-range gene flow across the sampled distribution of *P. alecto* and *P. conspicillatus*. Weights (*w*) on the edges that represent migration rates between pairs of demes are shown relative to their mean value (*<u>w</u>*) across all edges (Marcus et al., 2021). High migration regions are shown in blue and low migration regions in orange. Weakly supported long-range migration edges are indicated by dashed arrows, which are colored by the estimated source fraction (*ĉ*) (Shastry et al., 2025). C) Genetic differentiation is correlated with geographic distance in *P. alecto*. The first principal component (Table S11) is plotted against the geographic distance of each sample from Sawu, Indonesia, which is at one of the edges of the ring. D) Pairwise genetic distance in *P. alecto* is positively correlated with geographic distance. Each dot represents one non-redundant pair of the 86 sampled *P*. *alecto* individuals and the color gradient reflects the density of these dots. Distance reflects the number of allelic differences between two individuals as a ratio of the observed and the possible differences. A linear regression line was fitted to the data.

To further probe patterns of gene flow between distant populations, we used coalescent simulations to test multiple demographic scenarios including isolation and migration at three different time scales (Table S14). These coalescent simulations indicated recent migration after a period of isolation and divergence between *P. conspicillatus* populations and all *P. alecto* populations, forming a ring connecting Indonesia, Australia, and New Guinea (Table 1 and Figure S16B). Migration of *P. alecto* between Indonesia and Australia occurred recently but ceased in North West Australia and Burketown 1,380 ya (95% CI: 164–2,372 ya) and in the rest of Australia 4,016 ya (95% CI: 2,988–5,204 ya). The observed joint site frequency spectrum of our populations is recapitulated in simulations based on our inferred parameters, supporting a good model fit (Figure S17). Such migration in recent history is consistent with the empirical evidence of migration between Indonesia and Australia from the three historical *P. alecto* individuals collected in North West Australia from 1989 to 2003 which genetically clustered with Indonesian individuals (Figure 4A). However, the inferred cessation of gene flow between Indonesia and Australia suggests that this was a rare event.

**Table 1.** Inferred demographic parameters for *Pteropus alecto* and *P*. *conspicillatus* population pairs based on coalescent simulations. The best of four migration scenarios (no migration, ancient migration, recent migration, and constant migration) was selected based on the Akaike information criterion. Years since migration began are shown for recent migration and years since migration ended are shown for ancient migration. Estimates shown were selected from 100 replicates based on the likelihood. For additional inferred demographic parameters and 95% confidence intervals see Table S15. The following population codes are used: INDO+3NWA, Indonesia and three samples from North West Australia; NWA+BURK, remaining North West Australian samples and all Burketown samples; EA+NEA, East Australia and North East Australia; WTA+NG, Wet Tropics Australia and New Guinea. See Table S1 for population code assignments for each sample.

| Pop 1 | Pop 2 | n<br>(Pop 1) | n<br>(Pop 2) | Species comparison | Best<br>migration<br>scenario | Migration<br>from Pop 2<br>to Pop 1 | Migration<br>from Pop 1<br>to Pop 2 | Years since<br>migration<br>began/ended |
| --- | --- | --- | --- | --- | --- | --- | --- | --- |
| NWA+BURK | EA+NEA | 24 | 47 | alecto-alecto | ancient | $1.78 \times 10^{-4}$ | $5.99 \times 10^{-5}$ | 172 |
| INDO+3NWA | EA+NEA | 15 | 47 | alecto-alecto | ancient | $2.94 \times 10^{-5}$ | $3.28 \times 10^{-5}$ | 4,016 |
| INDO+3NWA | NWA+BURK | 15 | 24 | alecto-alecto | ancient | $5.75 \times 10^{-5}$ | $4.12 \times 10^{-5}$ | 1,380 |
| EA+NEA | WTA+NG | 47 | 55 | alecto-conspicillatus | recent | $5.72 \times 10^{-5}$ | $4.01 \times 10^{-5}$ | 25,020 |
| NWA+BURK | WTA+NG | 24 | 55 | alecto-conspicillatus | recent | $7.01 \times 10^{-5}$ | $4.66 \times 10^{-5}$ | 14,848 |
| INDO+3NWA | WTA+NG | 15 | 55 | alecto-conspicillatus | recent | $1.42 \times 10^{-5}$ | $9.82 \times 10^{-6}$ | 35,856 |

We inferred broadly similar migration scenarios and parameters when we treated Australian samples as a single population (Table S15). However, in this analysis migration between Indonesia and Australia was constant and showed a poor fit of observations to simulated data, which was driven by the inclusion of the three North West Australian samples clustering with Indonesian samples (Figure S18). This suggests it is challenging to fit a demographic scenario explaining the admixture patterns specifically between Indonesia and North West Australia, without modeling North West Australia as a discrete population. Similarly, assuming an average mammalian mutation rate rather than a higher human-based mutation rate does not broadly change inferred migration scenarios, though as expected it increases the time since recent migrations began (Table S16). Using the average mammalian mutation rate, we inferred that migration of *P. alecto* between Indonesia and North West Australia ceased 4,492 ya (95% CI: 1,132–7,680 ya) while migration of *P. alecto* between Indonesia and the rest of Australia ceased 11,316 ya (95% CI: 7,484–15,700 ya). In addition, we inferred a recent, rather than an ancient, migration scenario between North West Australia and the rest of Australia. Based on the AIC, neither mutation rate consistently produced better-supported demographic models (Table S14).

Although we inferred migration connecting the chain of three *P. alecto* populations from Indonesia to East Australia, we also found that genetic distance increased with geographic distance in this species (Figures 4C and 4D, Table S17). When investigating pairwise comparisons of genetic and geographic distance for all *P. alecto* individuals (Figure 4D), we found a significant positive correlation (Mantel test, r = 0.26, *p* = 0.001). Consistent with our population structure analyses, fastsimcoal2 estimates also showed consistently higher migration from *P. conspicillatus* populations to *P. alecto* populations. Divergence-times between *P. conspicillatus* and *P. alecto* were estimated by fastsimcoal2 using a human mutation rate to be 129–186 kya (Table S15). Using an average mammalian rate increases this divergence time to 368–554 kya (Table S16). Bayesian species-level divergence-time inference using a calibration point from an earlier dated phylogeny suggested that *P. alecto* and *P. conspicillatus* diverged 0.32-1.02 mya (Figure S19).

## 4 Discussion

Our investigation of genome-wide SNPs across 242 flying-fox individuals from four species revealed weakly differentiated or panmictic Australian populations and strong isolation-by-distance in Indonesian *P. alecto*. In Australian *P*. *alecto*, we identified an east-west genetic split, with demographic modeling estimating that migration between the two populations ceased approximately 172 years ago. This pattern may reflect the geographic barrier posed by the Gulf of Carpentaria, potentially reinforced by differences in phenology or mate preference between the two populations. Both *P. poliocephalus* and *P. scapulatus* show no evidence of population structure within Australia, consistent with their large populations (Macdonald et al., 2021; Vanderduys et al., 2024), higher mobility relative to *P. alecto* (Welbergen et al., 2020), and limited geographic barriers in their eastern Australian range. Interestingly, we found that the Indonesian *P. alecto alecto* taxon (Andersen, 1912) forms a distinct lineage to the other *P*. *alecto* samples, diverging from *P. alecto* and *P. conspicillatus* 2.61 mya. Our analyses suggest that *P. alecto alecto* is a previously unrecognized species, which should be considered in future taxonomic work on *Pteropus*.

We expected higher genetic differentiation between disjunct *P. conspicillatus* populations, instead finding limited evidence for isolation between these populations based on summary statistics and genetic structure analysis, suggesting that the roughly 150 km separating Australia and New Guinea do not represent a substantial geographic barrier for *P. conspicillatus*. Alternatively, the *P. conspicillatus* populations may be isolated but their divergence may be too recent to be apparent in our population genetic analyses. Other flying-fox species including *P*. *vampyrus* and *P*. *niger* have also been shown to have limited population genetic structure and high connectivity between populations (Larsen et al., 2014; Tsang et al., 2018), though species such as *P. dasymallus* are known to exhibit isolation-by-distance between island populations (Chen et al., 2021).

High levels of population connectivity across their ranges and large effective population sizes help explain the high genome-wide heterozygosity, an important measure of genetic diversity, which we found in all four species. Indeed, *P*. *alecto* is known to harbor higher genome-wide heterozygosity than other mammals (He et al., 2023; Zhang et al., 2013). In our study, the use of a single *P*. *alecto* reference genome for all species even increased the potential for underestimation of genetic diversity and divergence in the more diverged *P*. *scapulatus* and *P*. *poliocephalus*, though the modest decrease in alignment rates of 2%–3% for these species should only have a limited effect on our diversity estimates. A minor decrease in estimated diversity across species may also result from 6.5%–7.5% of per-species genome coverage targeting coding regions rather than the 3.8% expected under fully unbiased sampling, which likely reflects overrepresentation of GC-rich coding regions that can arise in ddRAD sequencing due to biases in enzymes and DNA fragment size selection (DaCosta and Sorenson, 2014). Despite this potential underestimation of genome-wide heterozygosity, when compared to genome-wide heterozygosity in a diverse set of mammals ranging from 0.012% in domestic cat to 0.16% in gorilla (Cho et al., 2013), our findings of genome-wide heterozygosity of 0.37% in *P*. *alecto* and 0.56% in *P. scapulatus* suggest that flying-foxes may generally be outliers with higher genetic diversity than other mammals.

Overall, we found that the higher degree of genetic diversity in Australian flying-fox populations compared with other populations is not primarily a result of admixture with sympatric species, as has been suggested (Tsang, 2015), but of strong within-species gene flow across Australia. Nevertheless, we found evidence of moderate hybridization between *P*. *alecto* and *P. conspicillatus* (Almeida et al., 2014). Consistent with hybridization rather than incomplete lineage sorting, admixed individuals primarily occurred in the broad region of distribution overlap in eastern Australia. However, hybrid backcross *P*. *alecto* individuals were not restricted to the narrow secondary contact zone with *P. conspicillatus* in the Wet Tropics region of Queensland, Australia. Instead, consistent with the high mobility of flying-foxes, hybrid backcrosses were widely distributed across eastern and northern Australia. We did not find any F_1_ hybrids, and the modest number of substantially admixed individuals detected suggests that hybridization is ongoing but infrequent.

Field observations indicate that the morphologically highly similar *P*. *alecto* and *P. conspicillatus* inhabit ecologically distinct niches, with *P. alecto* found in savanna woodland and rainforest (Palmer et al., 2000) and *P. conspicillatus* primarily in rainforest (Richards, 1990). Although *P. conspicillatus* is not a true rainforest specialist (Parsons et al., 2007), this general divergence in ecological specialization may contribute to limiting gene flow between both species. This ecological differentiation may also have played a role in the initial divergence of these two species since niche conservatism can isolate diverging populations in distinct ranges during early allopatric speciation, but does not prevent secondary contact during later range shifts (Lin et al., 2025). Differences in habitat choice may also contribute to the asymmetric gene flow, which is substantially higher from *P. conspicillatus* to *P. alecto*. If *P*. *alecto* is spreading into rainforest habitats occupied by *P*. *conspicillatus*, then such a range expansion can lead to asymmetric introgression of local genes into the genome of the encroaching species (Excoffier et al., 2009). Whether such asymmetric gene flow is contributing to local adaptation via introgression of beneficial alleles could be investigated in future studies using whole genome sequencing. Alternatively, asymmetric gene flow can also occur due to asymmetric mating between the sexes, as was found in two closely related horseshoe bat species (Mao et al., 2013). While *P*. *alecto* and *P. conspicillatus* do not differ substantially in size or morphology (Neaves et al., 2018), asymmetric mating could still occur due to differences in mating behavior and timing.

We did not find substantial evidence supporting our hypothesis that *P. alecto* and *P. conspicillatus* form an interbreeding population with gradual isolation-by-distance along the geographic ring connecting Australia, Indonesia, and New Guinea. Our findings that Australian populations of the two species interbreed and Indonesian *P. alecto* show strongest genetic differentiation from *P. conspicillatus* superficially supports the hypothesis of a ring species. Although the gene flow we infer between Australia and Indonesia depends on flying-foxes making a challenging journey of around 500 km in the present day, bats may use vegetated islands (such as those at Ashmore reef) and oil rigs as stepping stones. Historically, dispersal distances during the Pleistocene have also been substantially lower as the Sahul Shelf extending from Australia was exposed during periods of lower sea levels (Karin et al., 2020; Voris, 2000). However, we did not sufficiently sample the hypothesized endpoint of the ring connecting Indonesia to New Guinea to confirm that admixture ceased in this region. Additionally, the substantial divergence between the species on the Australian mainland, despite limited geographic barriers for such mobile animals, suggests that *P. alecto* and *P. conspicillatus* may have diverged in allopatry on the islands of the Indo-Australian Archipelago and later colonized Australia, establishing secondary contact and admixing. Alternatively, divergence could have proceeded through sympatric or parapatric speciation driven by ecological specialization, behavioral isolation, or reproductive barriers accumulating in genomic islands (Ravinet et al., 2017; Richards et al., 2019). Such scenarios of speciation and secondary contact are also supported by our demographic analysis. Unfortunately, the resolution of biogeographic analyses in flying-foxes is limited and only indicates *P. alecto* and *P. conspicillatus* originated in the broad region of Australia, New Guinea, and the South Pacific (Tsang et al., 2019). Based on Bayesian divergence-time estimates, we found *P. alecto* and *P. conspicillatus* diverged in the Pleistocene as recently as 0.32–1.02 mya, which was consistent with demographic modeling estimates of 0.13–0.55 mya. This time was shaped by increases in land area and elevation in Wallacea (Hall, 2013) as well as the uplift of New Guinea ca. 5 mya, which played a key role as an ecological “stepping stone” between Wallacea and the Australian mainland (Roycroft et al., 2022). Notably, Australia and New Guinea were connected at times of low sea level during the Pleistocene including up to ca. 8 kya (Dodson, 1989; Voris, 2000). During the Holocene, human-wildlife conflict (Preece, 2023) and the reduction of rainforest habitat in Australia (Rule et al., 2012) may also have substantially impacted flying-fox distributions. These geographic upheavals provided ample opportunities for speciation, but also complicate reconstruction of flying-fox biogeography to establish the existence of a ring species. Moreover, confirming a ring species requires strong evidence since seemingly clear cases of ring species have been shown to result from more complex scenarios of vicariance, long-distance colonization, and allopatry (Liebers et al., 2004). Extended sampling of the range of *P. alecto* and *P. conspicillatus* outside of Australia combined with morphological analysis and whole genome sequencing could shed further light on the evolutionary origin of these species.

Our findings also have implications for conservation management of flying-foxes and pathogen control. *P. conspicillatus* is classified as endangered and hybridization, such as the cases we identified with *P*. *alecto*, may increase extinction risk (Todesco et al., 2016). However, the asymmetry of the gene flow between the two species, resulting in limited admixture in *P. conspicillatus*, suggests that there is limited increased extinction risk due to hybridization. Furthermore, the genetic connectivity between New Guinea and Australian *P. conspicillatus* populations may increase the resilience of the species by maintaining diversity. Unlike *P. conspicillatus*, *P*. *alecto* is currently listed as “Least Concern” by the International Union for Conservation of Nature (IUCN) across its range. However, the taxon *P. alecto alecto* identified here may be at risk, particularly since Indonesian populations in Sulawesi have declined due to hunting pressure since the 1970s (Sheherazade & Tsang, 2015). Our population genetic findings of high connectivity between flying-fox populations, including across international borders, can also inform studies of pathogen transmission and the management of biosecurity risk (Breed et al., 2010). For example, *P. alecto* has been implicated as a reservoir for bat-borne pathogens including Hendra virus (Edson et al., 2019). The high migration rates we inferred between *P. alecto* populations suggest that emerging viral strains can rapidly spread across the wide distribution of its host. Although these risks are well-understood and mitigated between Australia and New Guinea, our study suggests that transmission between northern Australia and Indonesia is underappreciated. Finally, hybridization between flying-foxes has also been suggested to explain the wide distribution of Nipah virus through interspecies viral transmission (Olival et al., 2020), implying pathogens hosted by *P. alecto* may readily be transmitted and spread by *P. conspicillatus*.

## 5 Conclusion

We have used genomic markers to provide new insights into the evolutionary history and population dynamics of flying-foxes, revealing high genetic connectivity across Australian populations and across disjunct *P. conspicillatus* populations in Australia and New Guinea. The isolation-by-distance observed in Indonesian *P. alecto* contrasts with the weak differentiation across large distances in other populations, highlighting the role of water barriers in shaping genetic structure. While our results do not support a ring species model for *P. alecto* and *P. conspicillatus*, they suggest a complex history of isolation-by-distance and admixture contributing to the evolution of these recently diverged species.

## 6 Data and code accessibility statement

Raw sequence reads and metadata are deposited in the SRA (BioProject PRJNA1230740). All code and genotype data used for the population genetic analysis and generation of figures is available via github (https://github.com/ascheben/pteropus_popgen_analysis).

## 7 Ethics approval statement

The animals sampled in this study were captured in accordance with animal ethics approval (Permit number AEC 2016-17).

## 8 Author contributions

**AS**: formal analysis, data curation, writing - original draft. **AMcK**: resources, data curation, validation, writing - review and editing. **TW**: resources, data curation, validation, writing - review and editing. **DAW**: conceptualisation, methodology, validation, writing - review and editing, supervision. **SSM**: investigation, writing - review and editing. **EPV**: resources, validation, writing - review and editing. **BLW**: conceptualisation, methodology, validation, writing - review and editing, supervision, funding acquisition.

## 9 Acknowledgements

This work was funded by the CSIRO Biodiversity, Ecosystem Knowledge and Services Program. We thank Owain Edwards for intellectual input and Samia Elfekih and Ian Cresswell for logistical support. We thank Tolga Bat Hospital, Bat Care Capricornia, Kimberly Wildlife Carers, Western Australian Museum, Port Moresby Botanical Gardens, National Flying-fox monitoring program (NFFMP), Cecelia Sanchez, Chris Todd, and Wayne Boardman for assistance in collecting samples. We also thank Christina DeDora and Jesse Ingram for manuscript editing support. Finally, we thank two anonymous reviewers for constructive comments.

## 11 Supplementary figures

**Figure S1.**
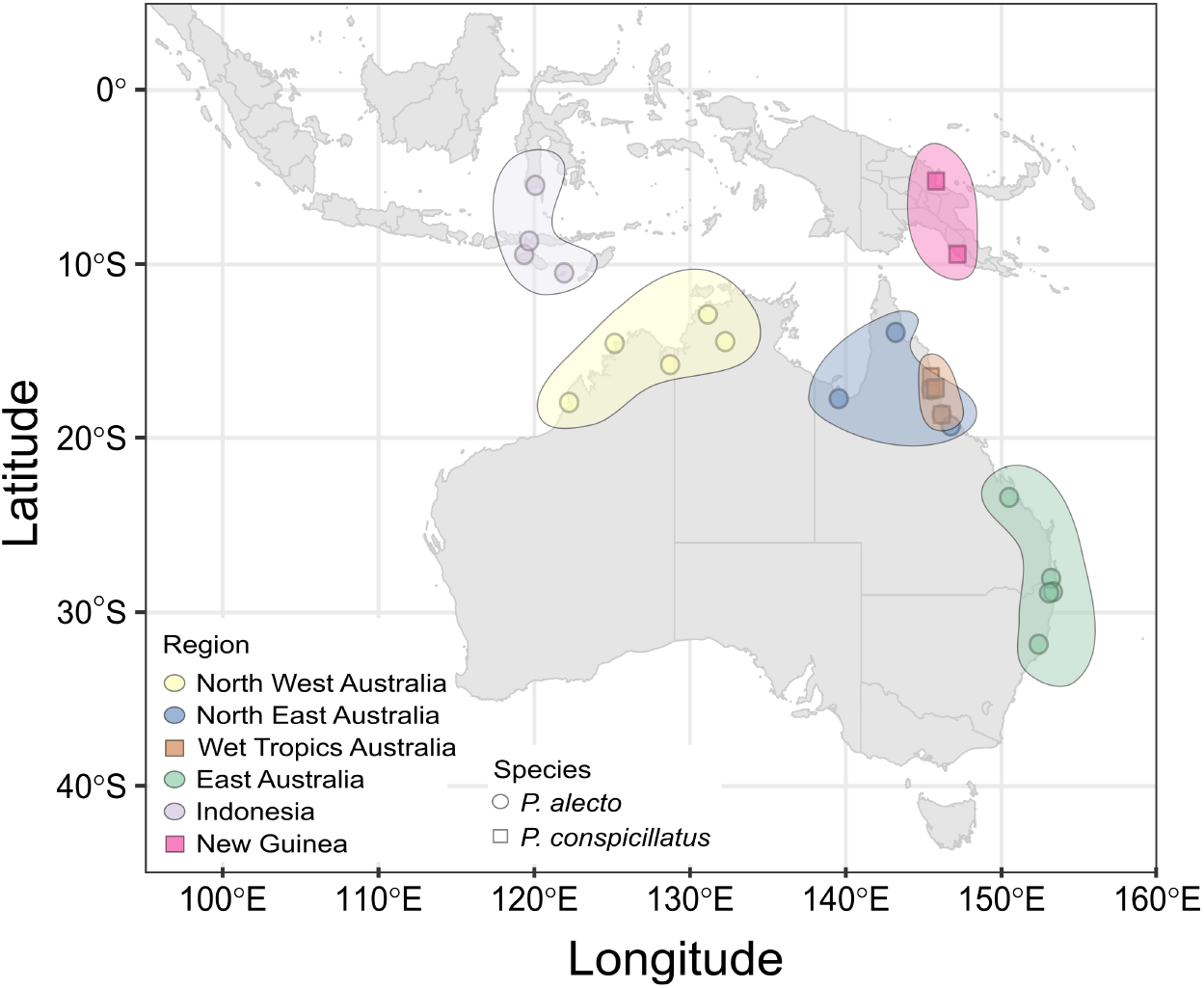
Roost locations and their assignment to distinct geographic regions sampled in this study. To highlight the regions relevant to this study, sampling locations here are shown only for the most widely distributed species *Pteropus alecto* and *P*. *conspicillatus*.

**Figure S2.**
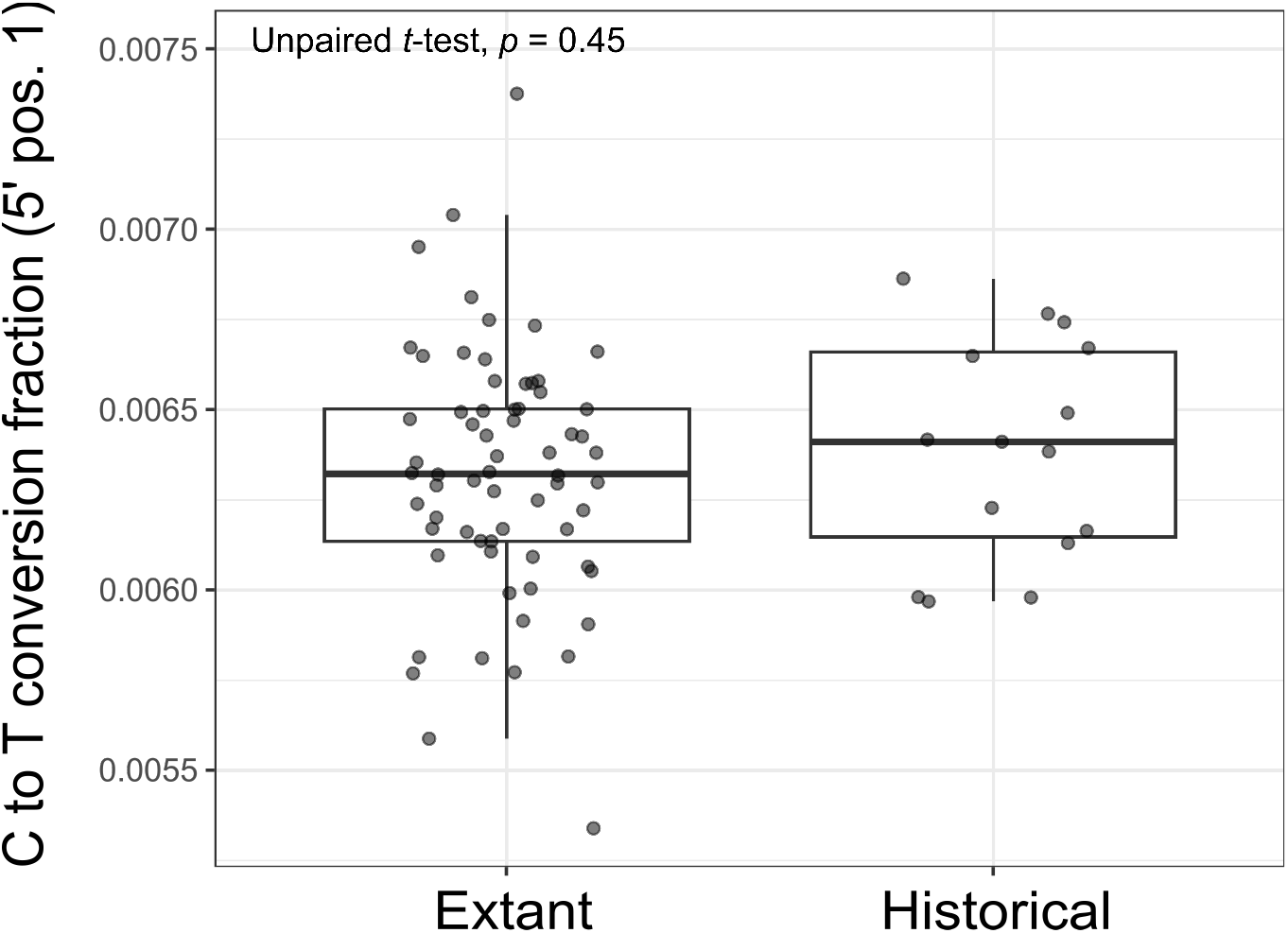
Comparison of flying-fox DNA degradation based on cytosine deamination between extant samples from CSIRO (n=66) and historical samples from the WA museum (n=15).

**Figure S3.**
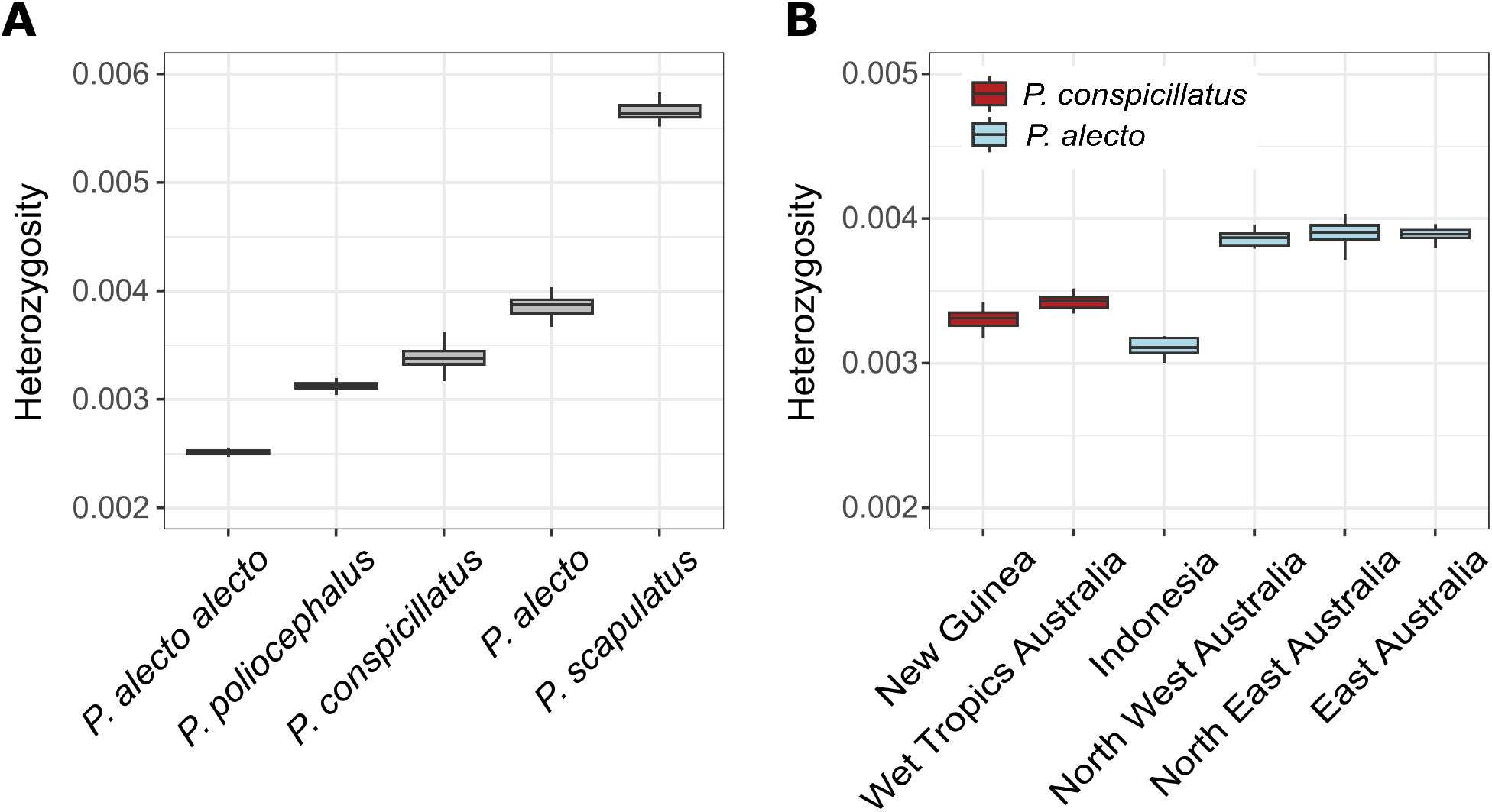
Genome-wide heterozygosity estimates for A) four flying-fox species and B) six regional populations of *Pteropus alecto* and *P. conspicillatus*. A total of 2,024,709 sites were used for *Pteropus alecto*, 2,518,218 for *P. conspicillatus*, 2,183,168 for *P*. *alecto alecto*, 2,610,770 for *P*. *poliocephalus*, and 2,409,166 for *P*. *scapulatus*.

**Figure S4.**
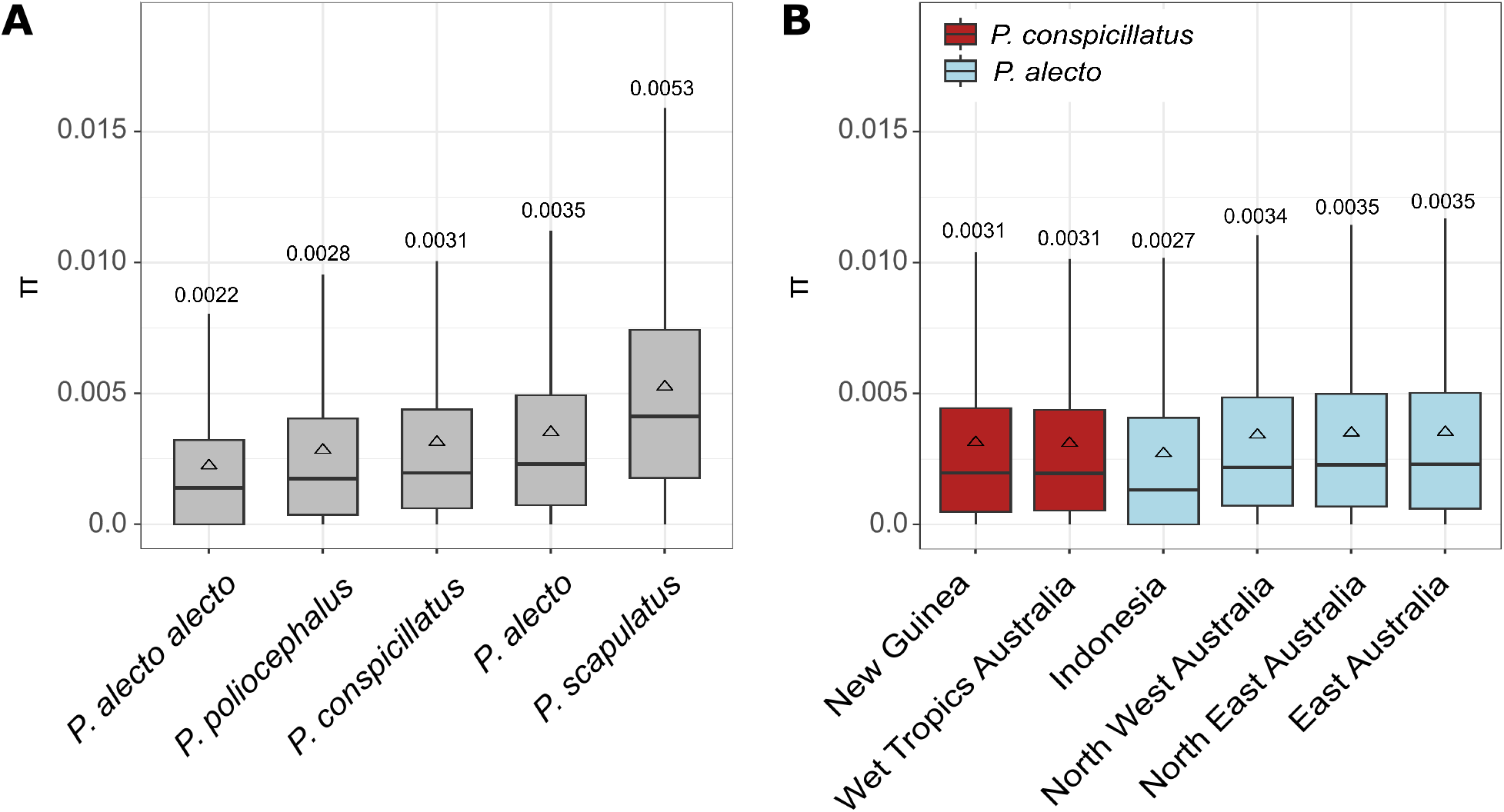
Genome-wide nucleotide diversity (*π*) estimates for A) four flying-fox species and B) six regional populations of *Pteropus alecto* and *P. conspicillatus*. Means are shown as triangles and specific mean values are shown above each boxplot. The number of sites used for each species is the same as listed in the Figure S3 legend.

**Figure S5.**
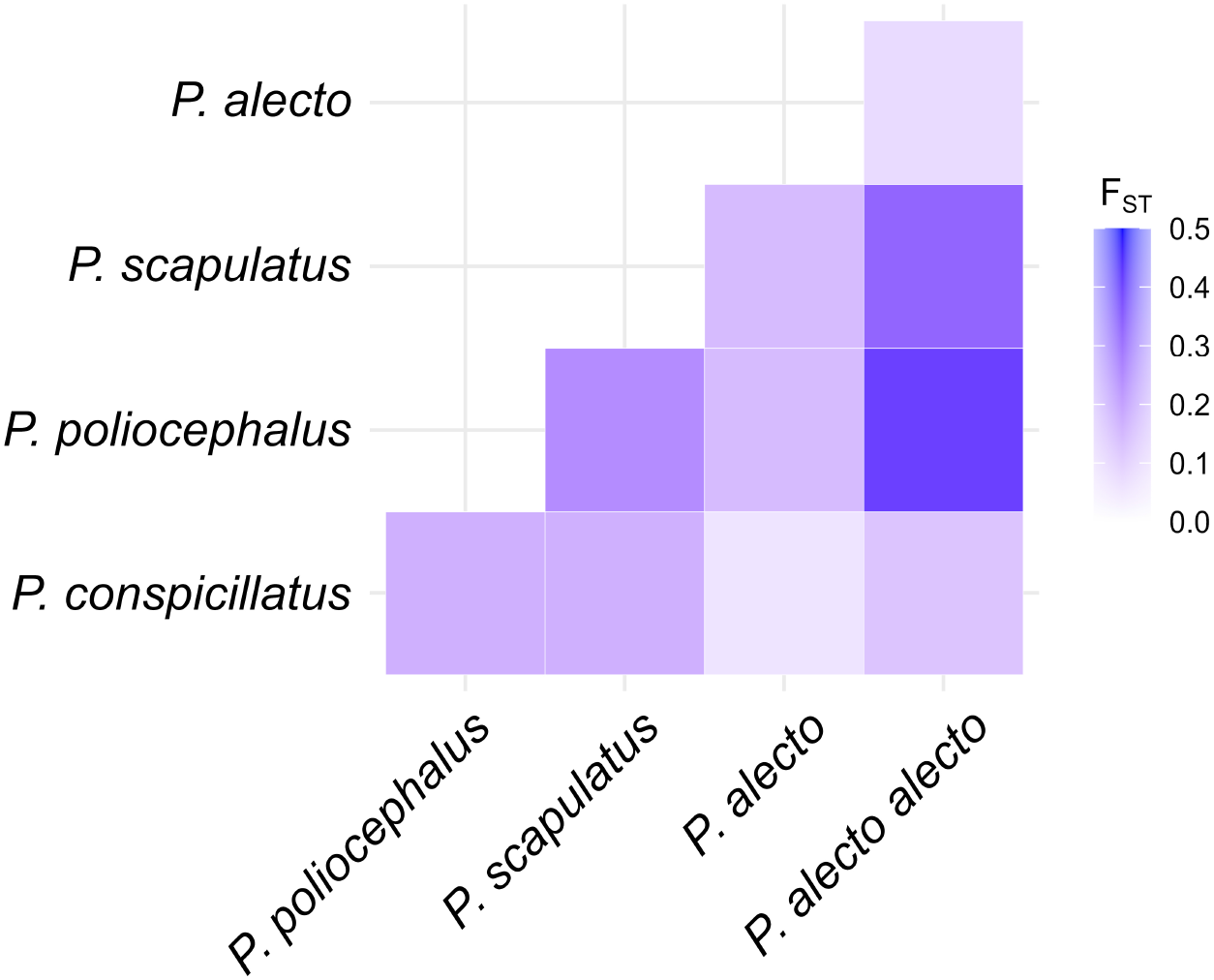
Genetic differentiation between four flying-fox species based on F_ST_.

**Figure S6.**
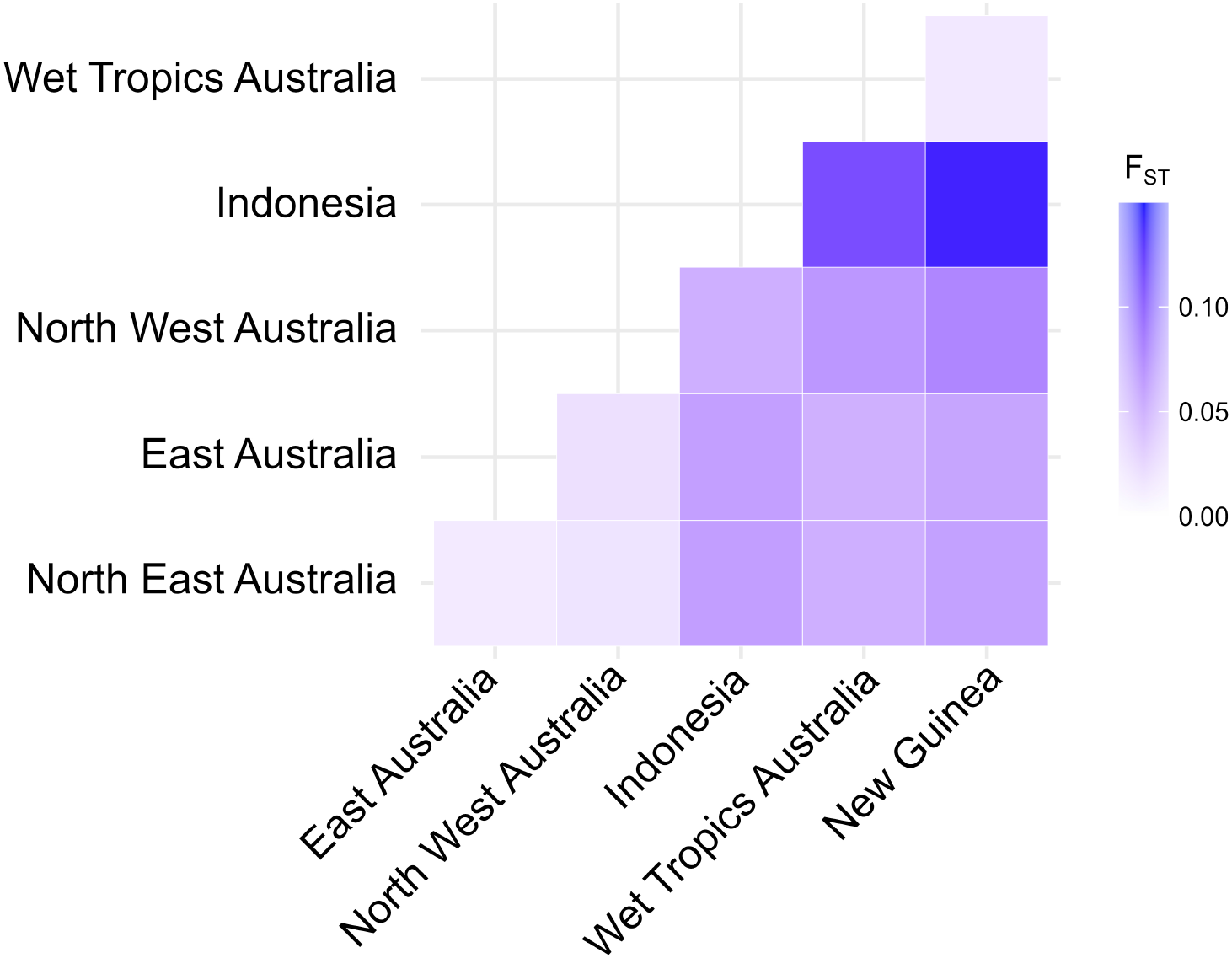
Genetic differentiation between geographic populations of *P*. *alecto* and *P*. *conspicillatus* based on F_ST_. The Wet Tropics Australia and New Guinea groups represent *P*. *conspicillatus* and the other groups represent *P. alecto*.

**Figure S7.**
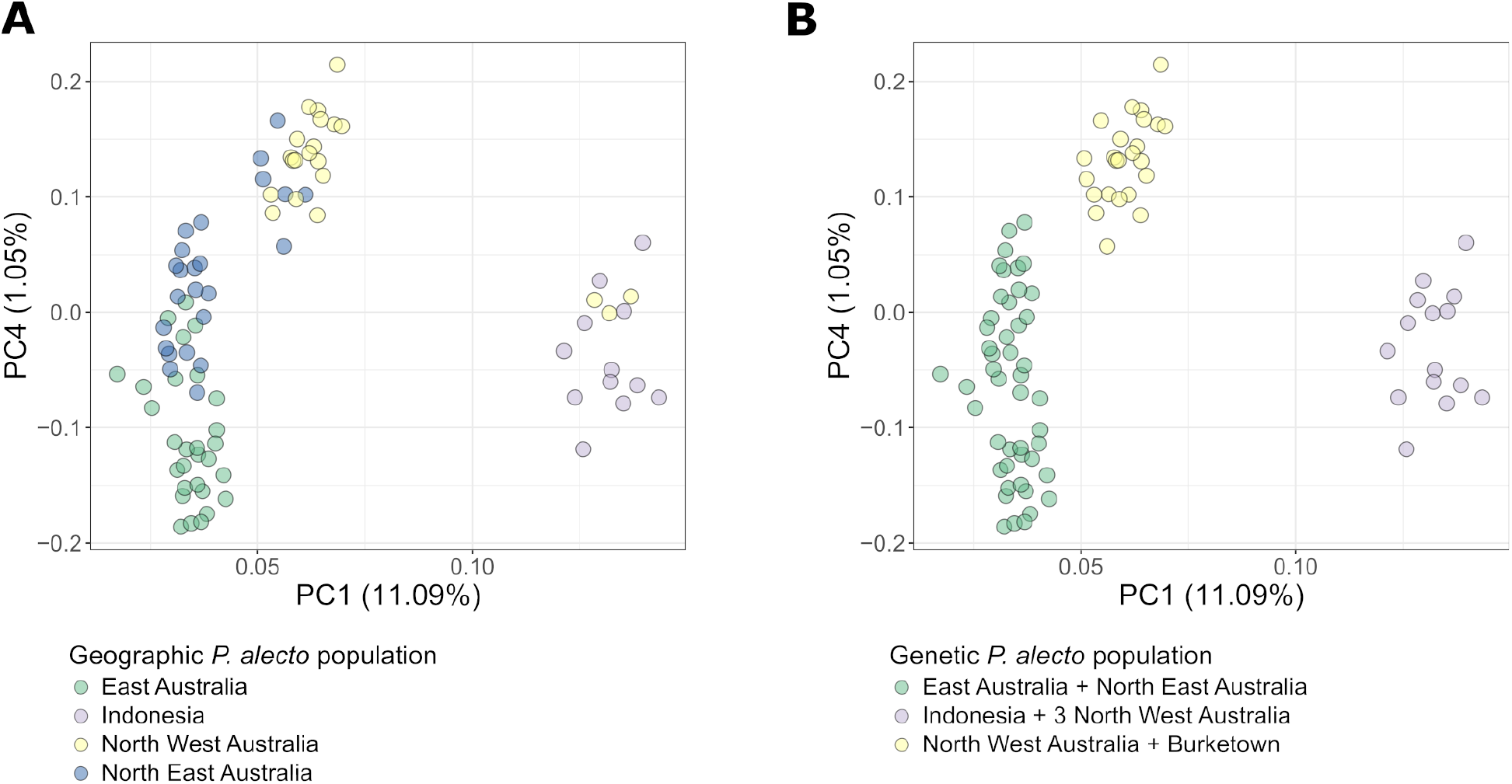
SNP-based principal component analysis (PCA) of *P. alecto* using the “alecto + conspicillatus” dataset colored by A) geographic populations and B) genetic populations.

**Figure S8.**
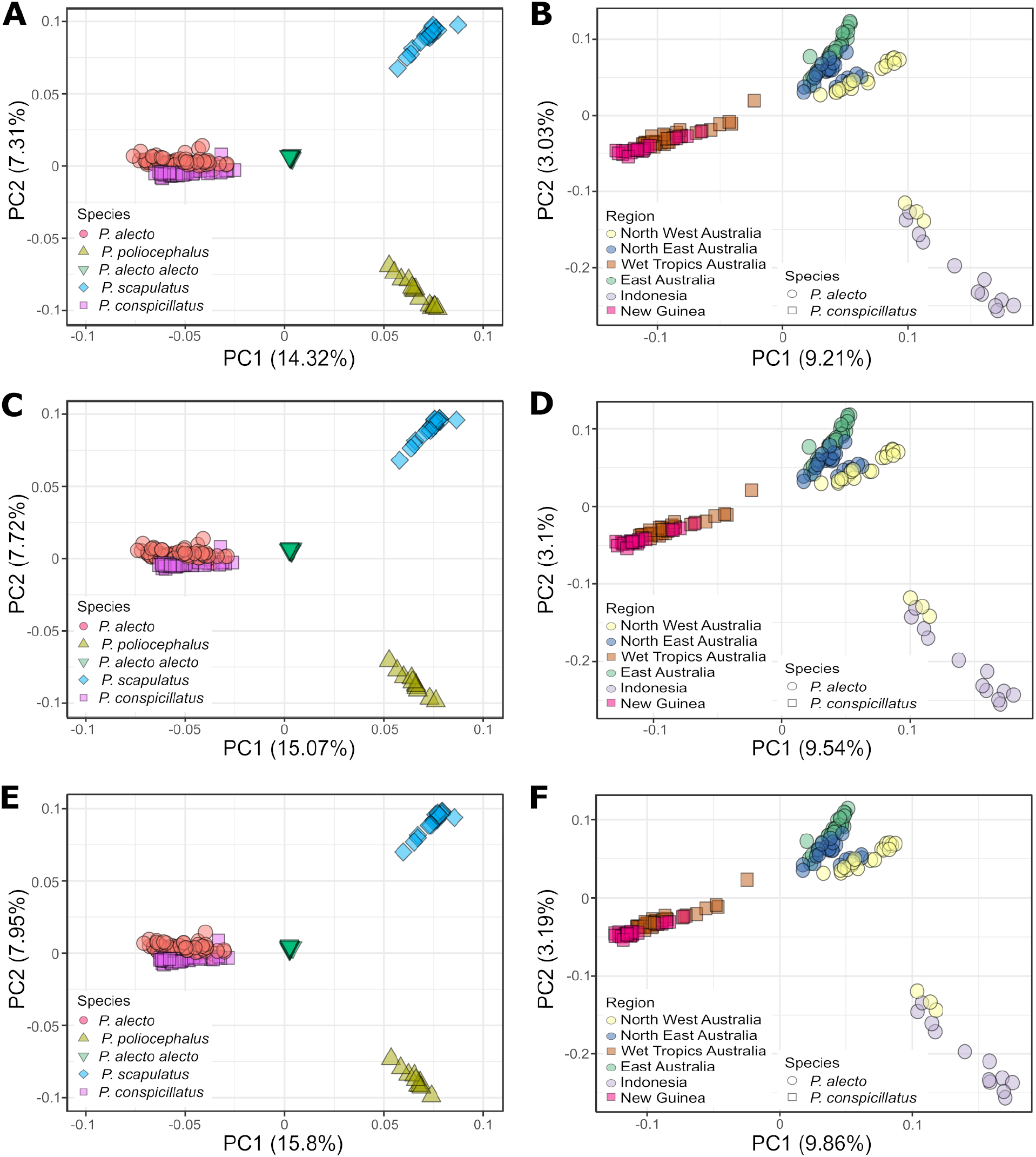
SNP-based principal component analysis (PCA) of flying-fox populations across three genotype missingness thresholds. For the species-level PCA, a genotype missingness threshold of 50% (22,394 SNPs) is shown in A), 40% (19,501 SNPs) in C), and 30% (16,160 SNPs) in E). For the population-level PCA of *P*. *alecto* and *P*. *conspicillatus*, a genotype missingness threshold of 50% (33,830 SNPs) is shown in B), 40% (29,222 SNPs) in D), and 30% (23,946 SNPs) in F).

**Figure S9.**
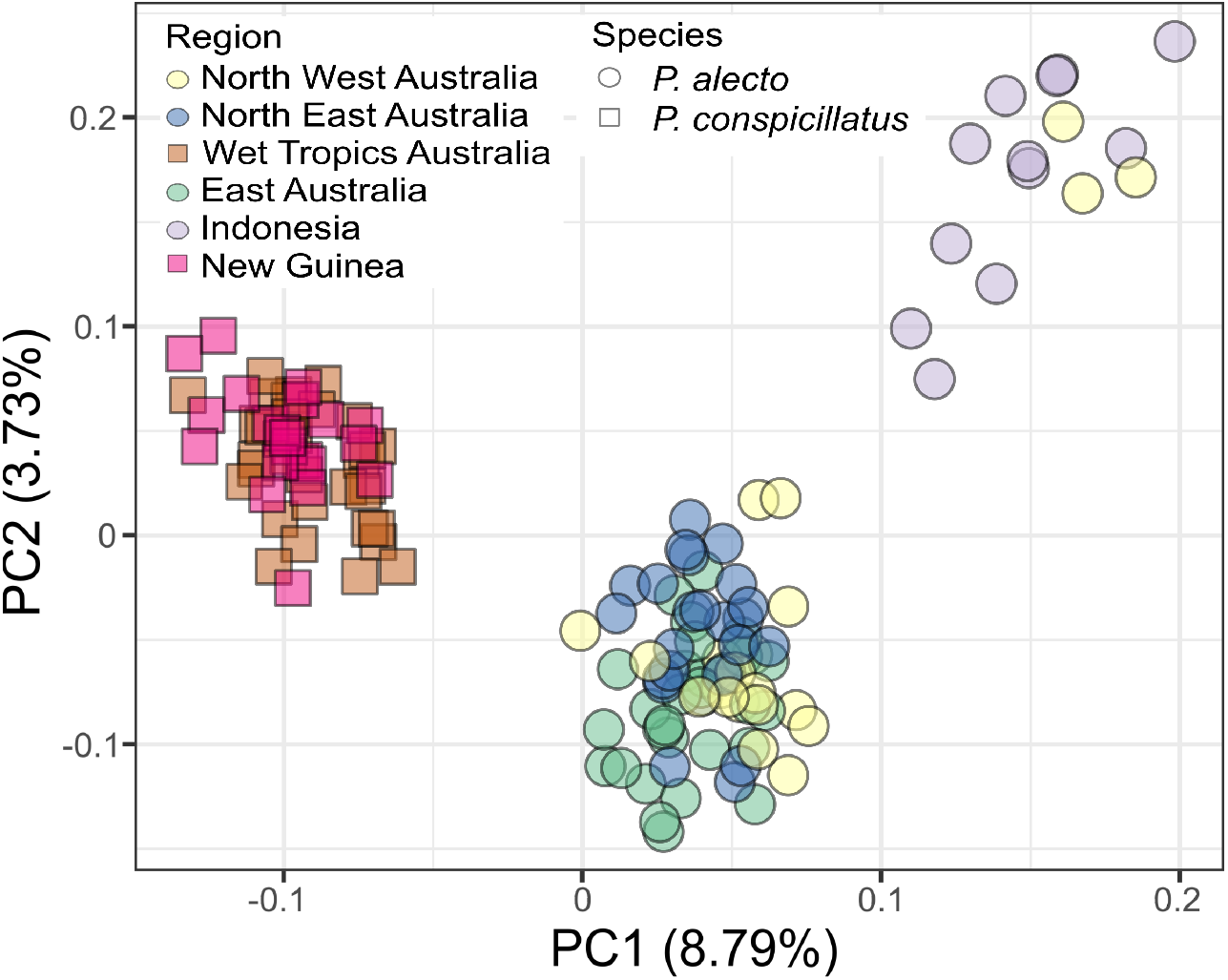
SNP-based principal component analysis (PCA) of *P*. *alecto* and *P*. *conspicillatus* populations using 216 SNPs with no missing genotypes.

**Figure S10.**
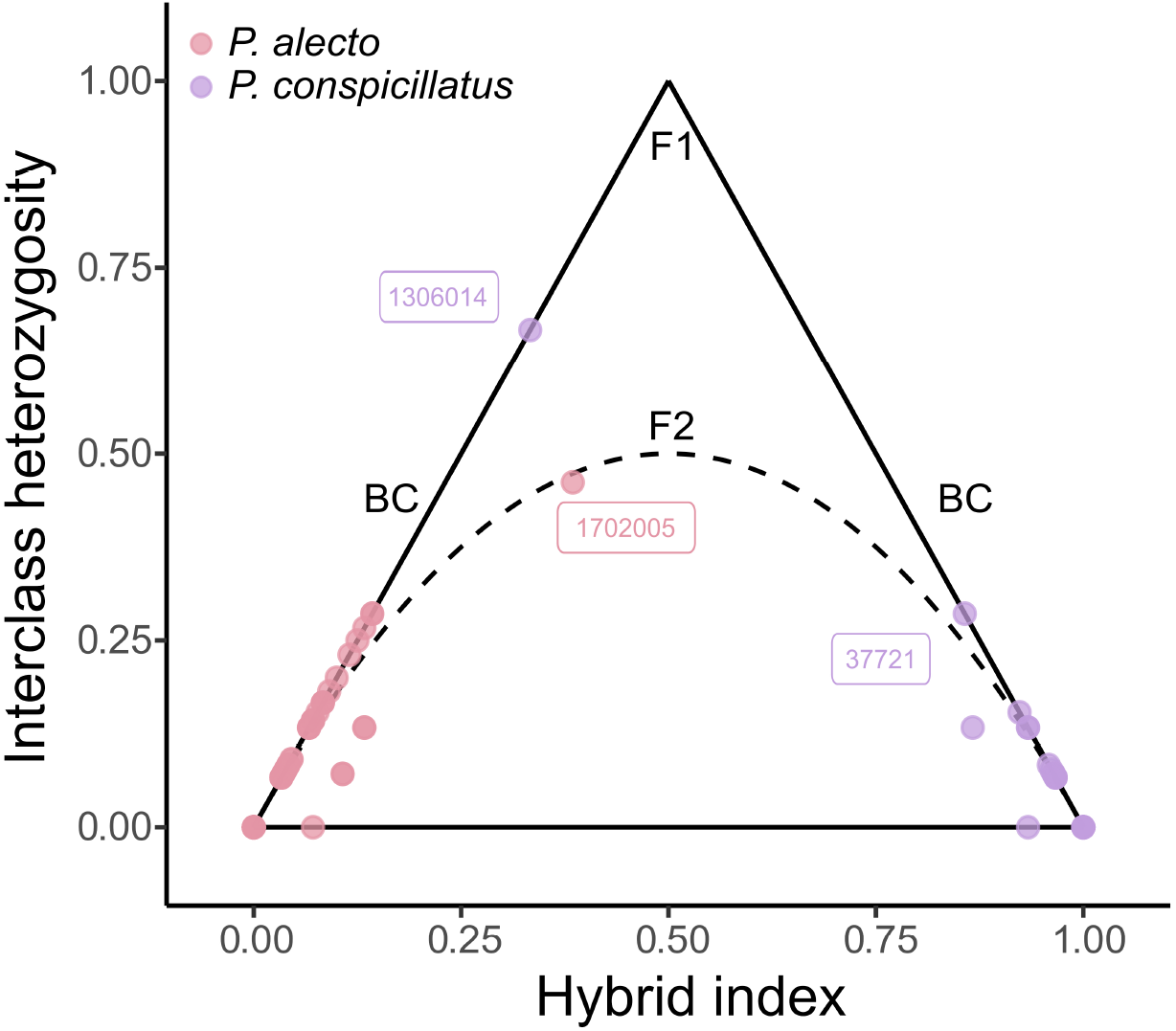
Triangle plot of admixture between *P*. *alecto* and *P*. *conspicillatus* individuals based on 15 ancestry-informative SNP markers. The hybrid index of F_1_ and F_2_ hybrids is expected to be 0.5 while backcross (BC) individuals between hybrids and parental lineages are expected to lie on the edges of the triangle with elevated heterozygosity and hybrid indices. Outliers are labelled with sample IDs in boxes. Proportions of missing SNP markers were not associated with high interclass heterozygosity or hybrid indices (Figure S11).

**Figure S11.**
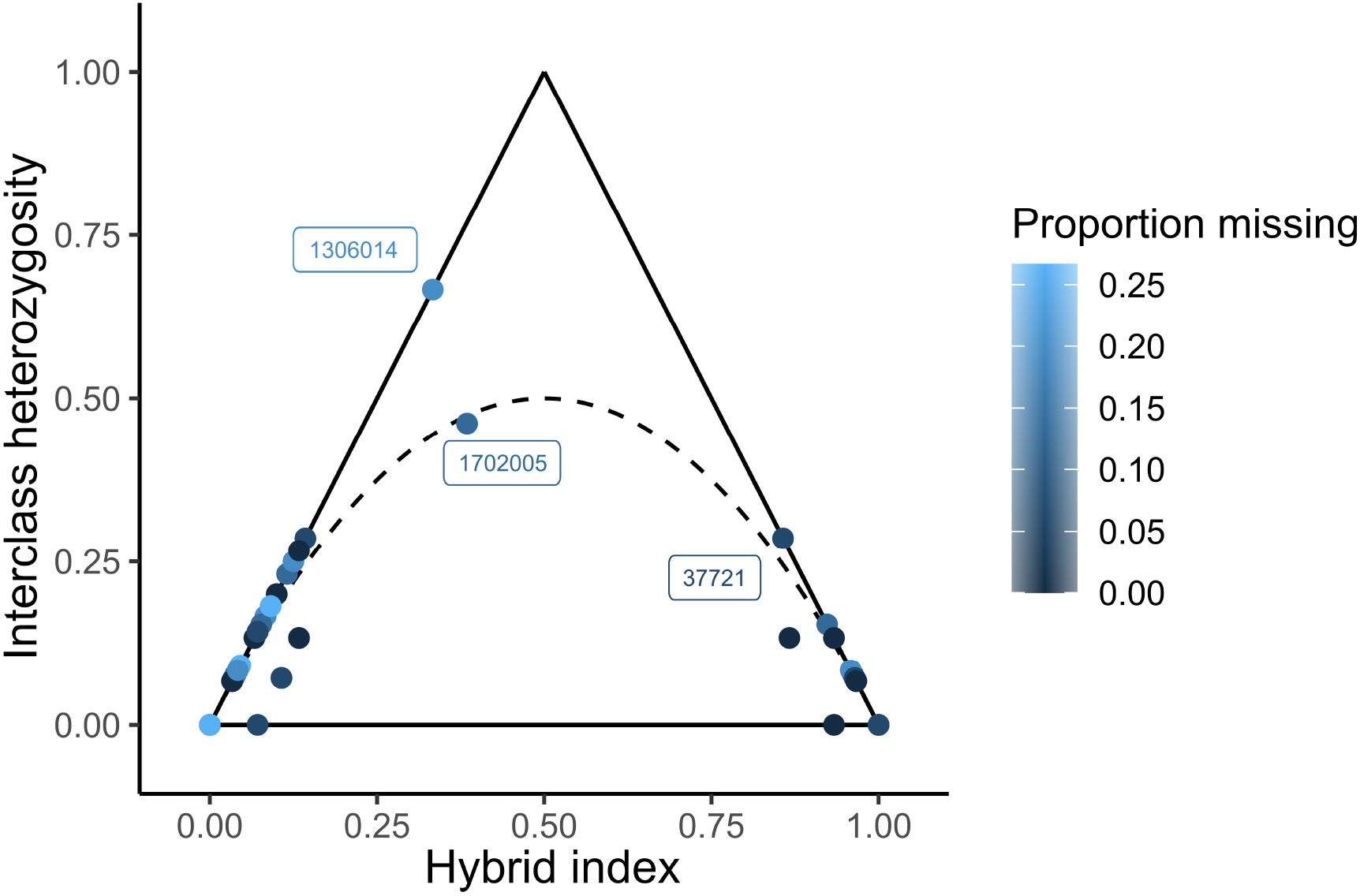
Triangle plot of admixture between *P*. *alecto* and *P*. *conspicillatus* individuals based on 15 ancestry-informative SNP markers colored by the proportion of missing markers per individual. Outliers are labelled with sample IDs in boxes.

**Figure S12.**
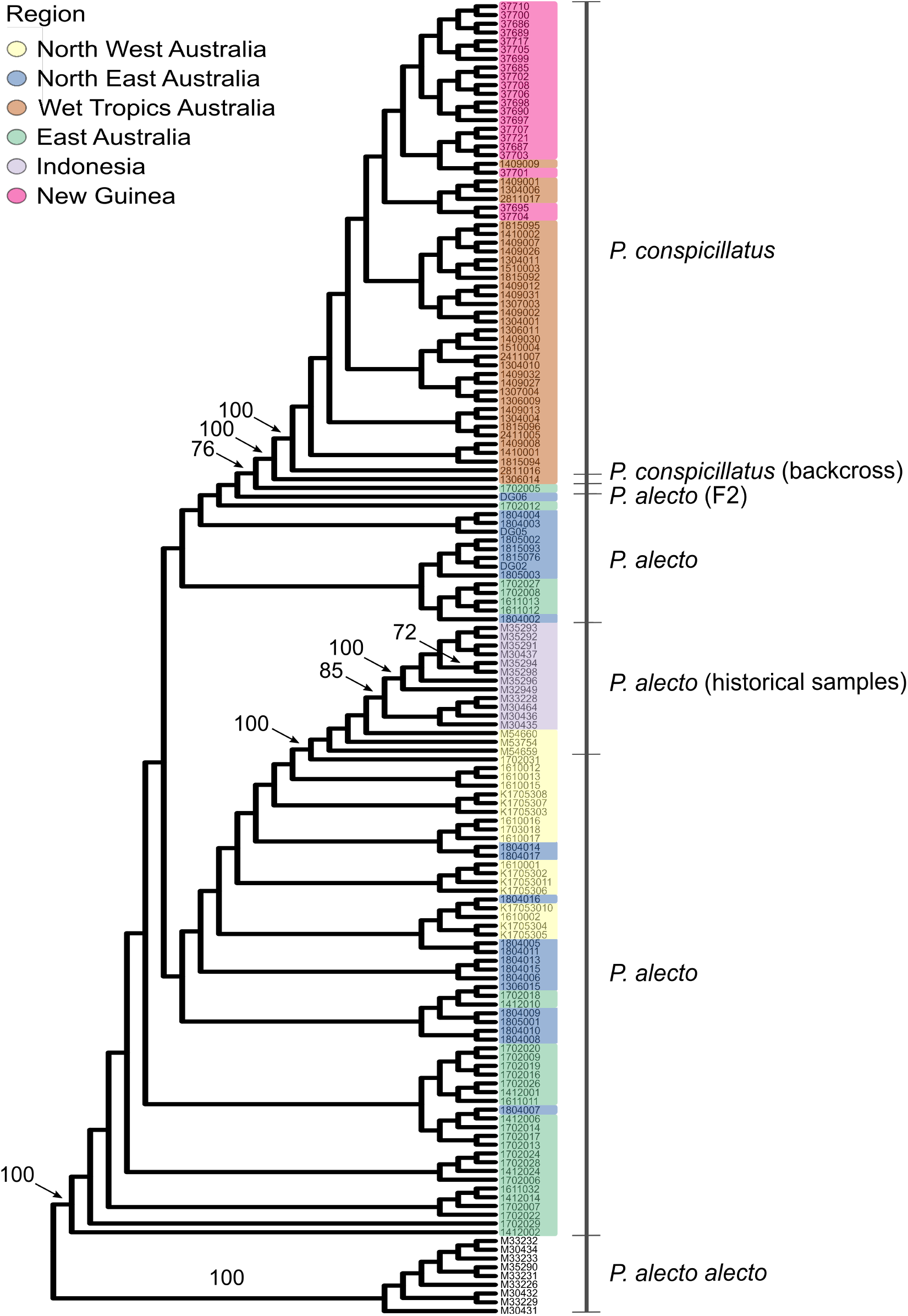
SNP-based maximum likelihood phylogeny of *Pteropus alecto* and *P*. *conspicillatus*. A total of 17,708 SNP sites were used. Individuals are highlighted in a color by geographic region and the Indonesian *P. alecto alecto* group is used as the outgroup. Historical *P. alecto* samples were collected from 1989 to 2003 (see Table S1). Bootstrap support values >70 are shown.

**Figure S13.**
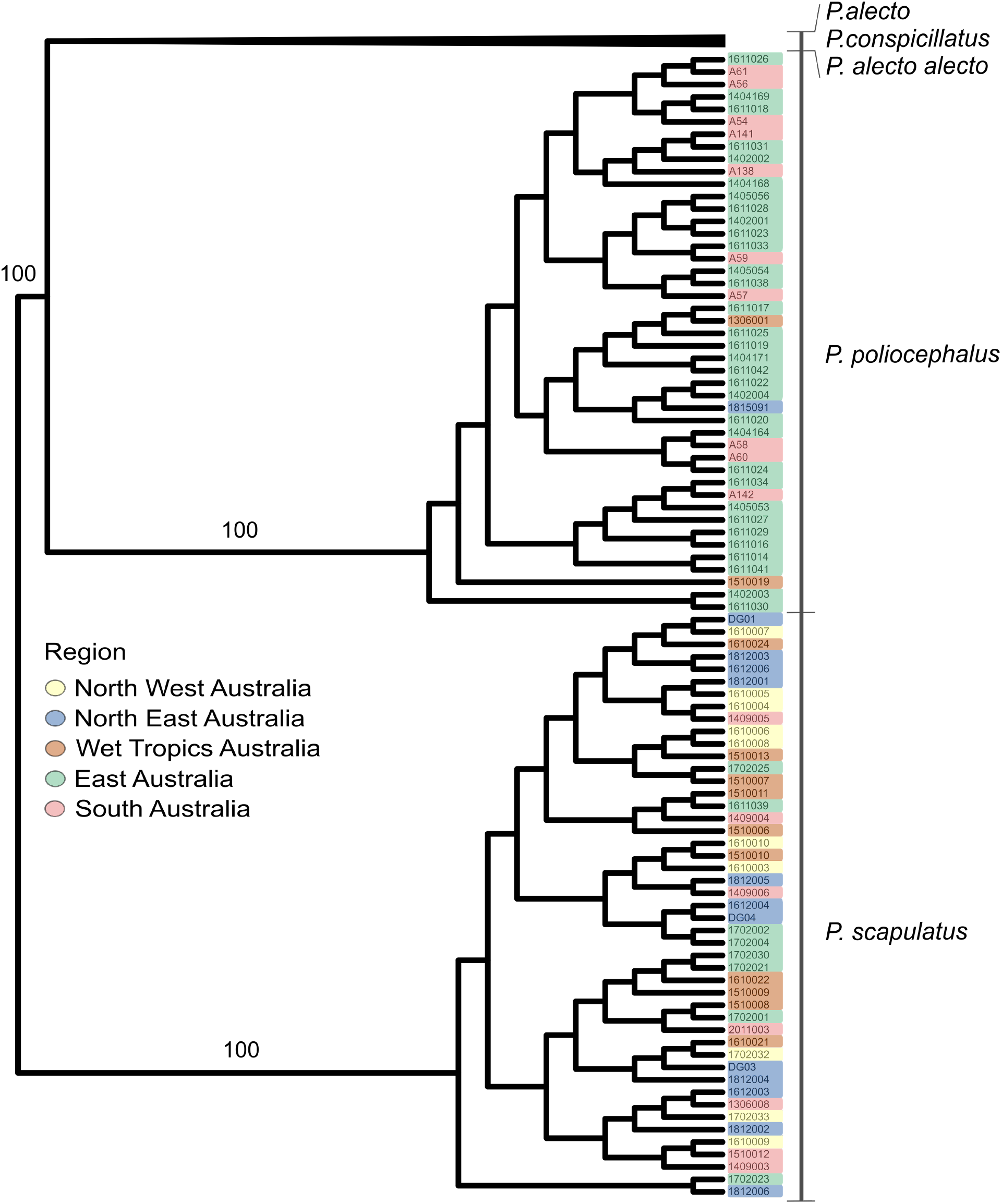
SNP-based maximum likelihood phylogeny of *Pteropus poliocephalus* and *P*. *scapulatus*. A total of 11,818 SNP sites were used. Individuals are highlighted in a color by geographic region. Bootstrap support values >70 are shown, highlighting robustly supported branches.

**Figure S14.**
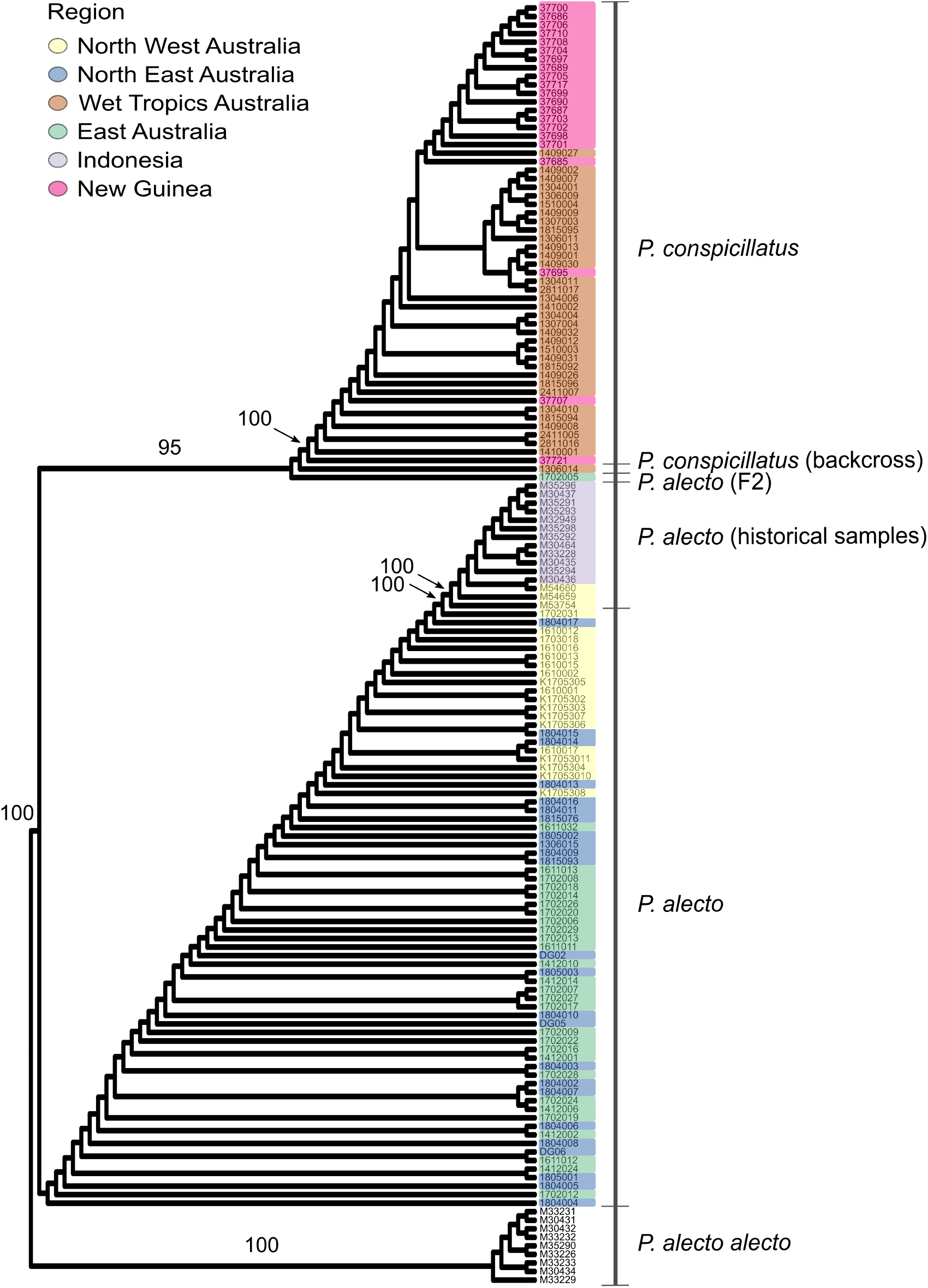
SNP-based phylogeny of *Pteropus alecto* and *P*. *conspicillatus* inferred by CASTER-site. A total of 17,708 SNP sites were used. Individuals are highlighted in a color by geographic region. Branch bootstrap support values >95 are shown.

**Figure S15.**
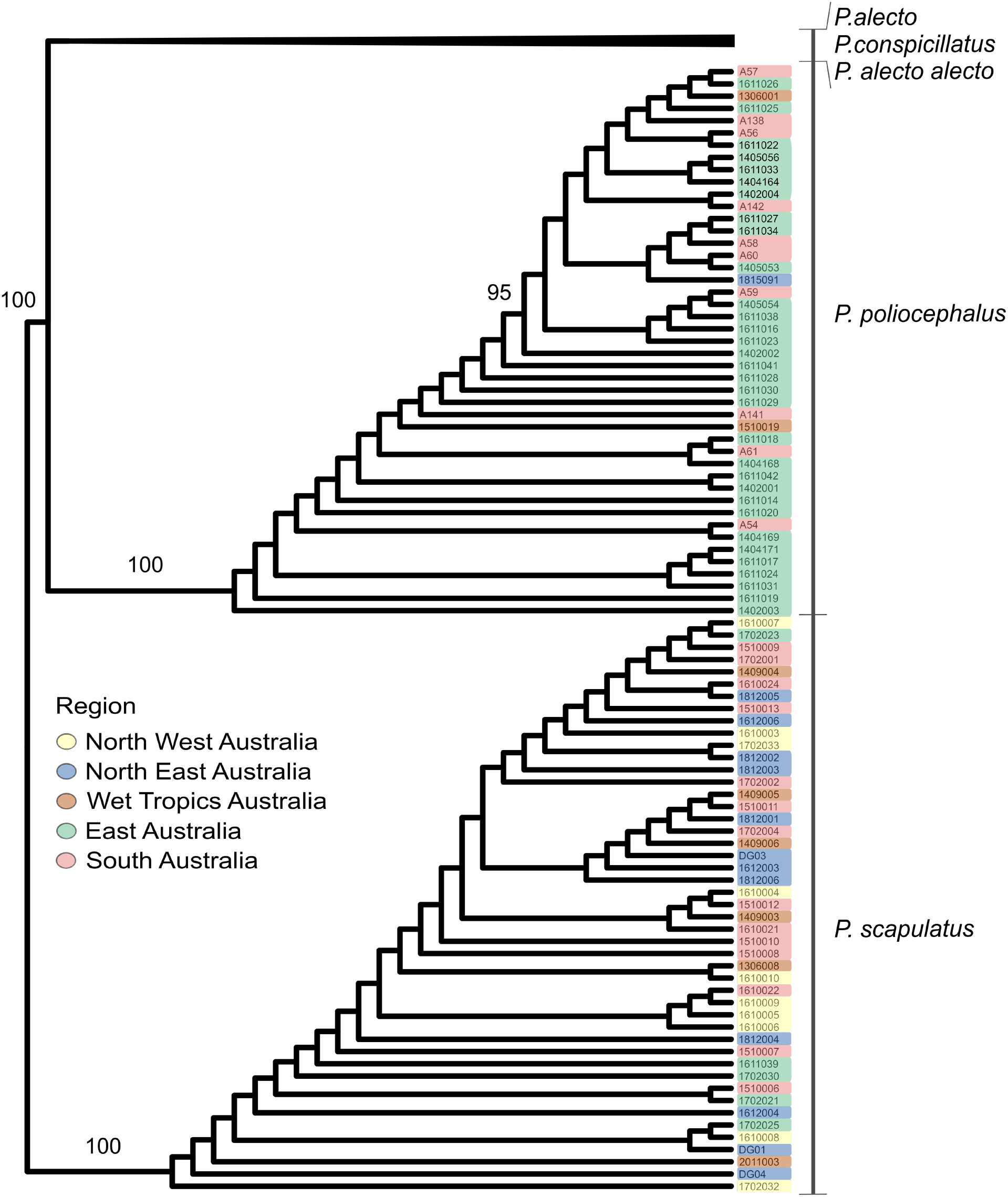
SNP-based phylogeny of *Pteropus poliocephalus* and *P*. *scapulatus* inferred by CASTER-site. A total of 11,818 SNP sites were used. Individuals are highlighted in a color by geographic region. Branch bootstrap support values >95 are shown.

**Figure S16.**
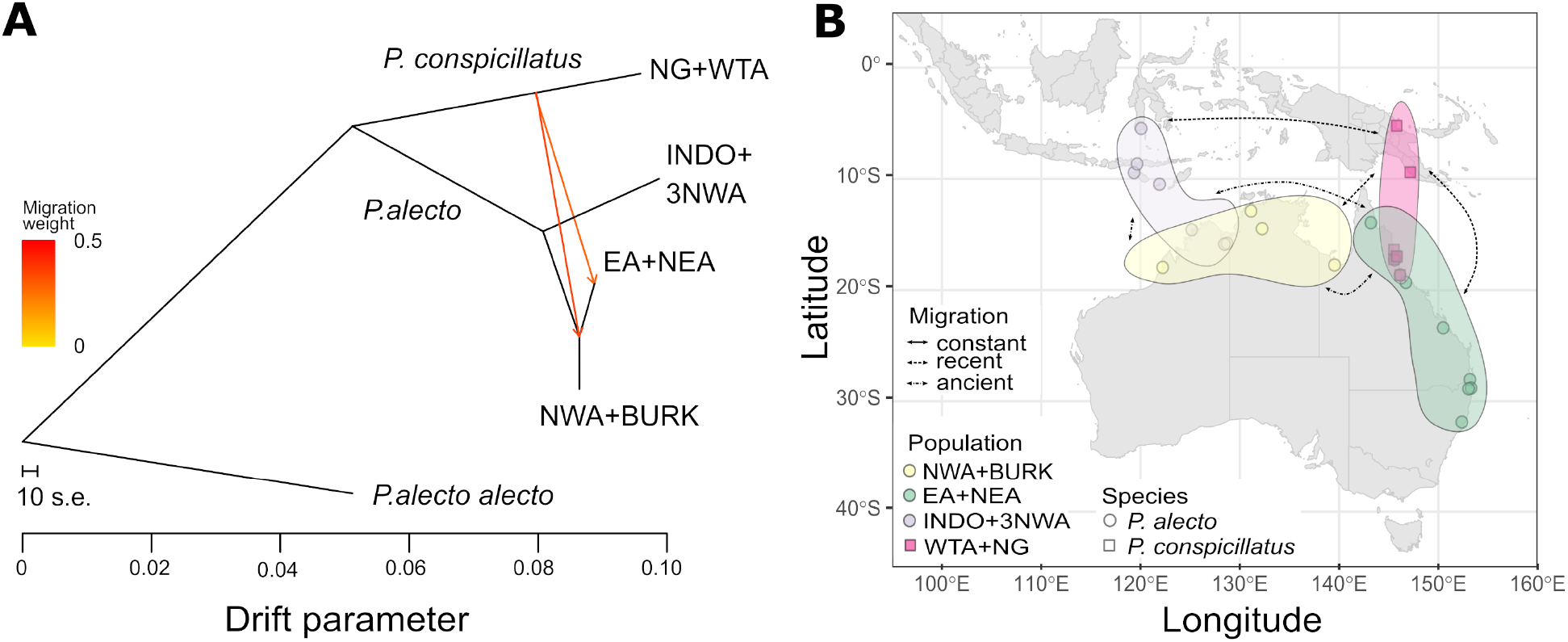
Gene flow in *Pteropus alecto* and *P. conspicillatus* along a geographic ring ranging across Indonesia, Australia and Papua New Guinea. A) Maximum likelihood tree with migration edges for three genetic *P. alecto* populations and *P. conspicillatus*. The diverged taxon *P. alecto alecto* was used as the outgroup. B) Map of inferred migration scenarios between populations using coalescent simulations (see Table 1 for additional inferred parameters). The following population codes were used in A) and B): INDO+3NWA, Indonesia and three samples from North West Australia; NWA+BURK, remaining North West Australian samples and all Burketown samples; EA+NEA, East Australia and North East Australia; WTA+NG, Wet Tropics Australia and New Guinea.

**Figure S17.**
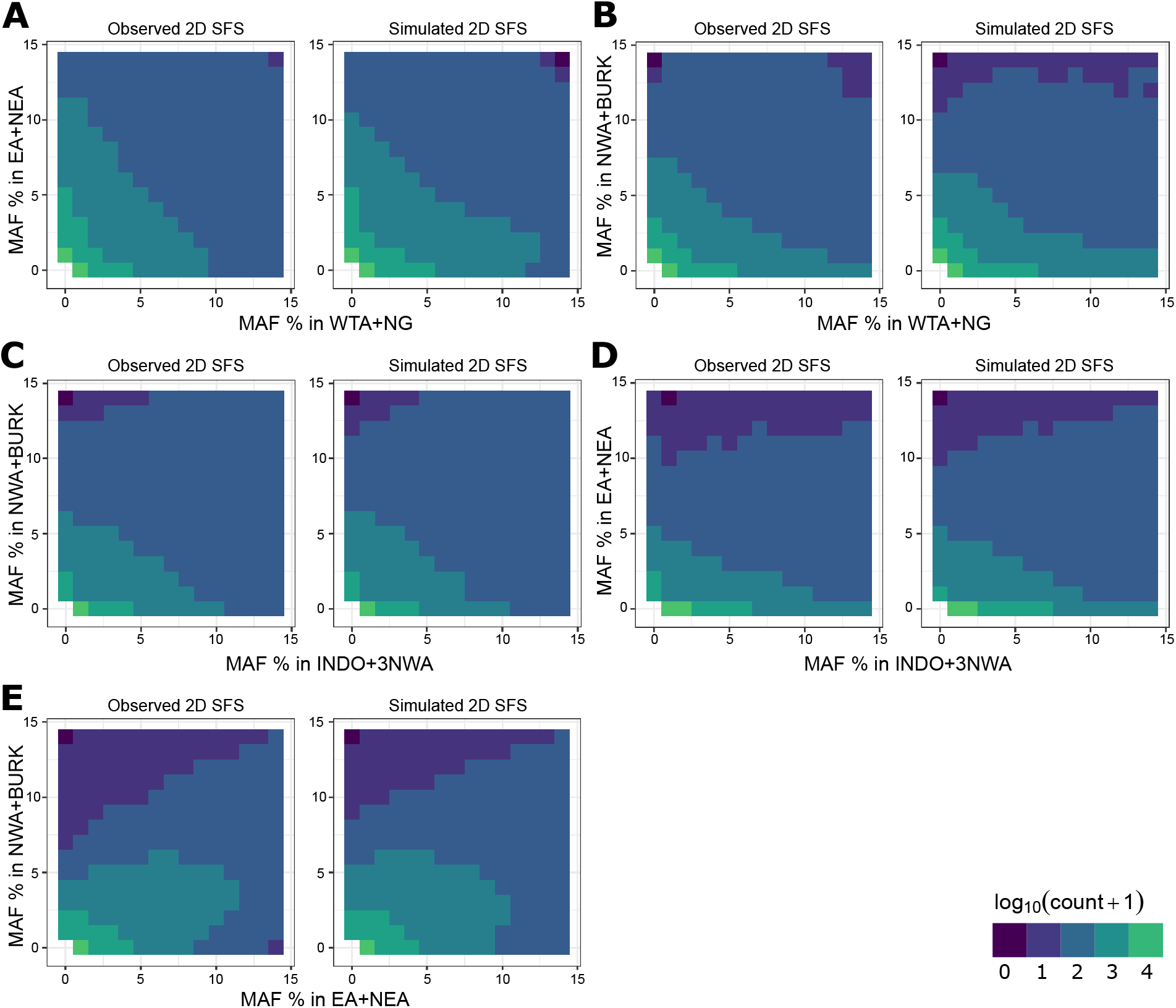
Comparison of the observed and simulated joint 2D site frequency spectrum (SFS) under the best-fit demographic model across flying-fox population pairs (A-E). Log-transformed counts of sites are shown for minor allele frequency (MAF) pairs. Monomorphic sites are omitted. The following population codes were used: INDO+3NWA, Indonesia and three samples from North West Australia; NWA+BURK, remaining North West Australian samples and all Burketown samples; EA+NEA, East Australia and North East Australia; WTA+NG, Wet Tropics Australia and New Guinea.

**Figure S18.**
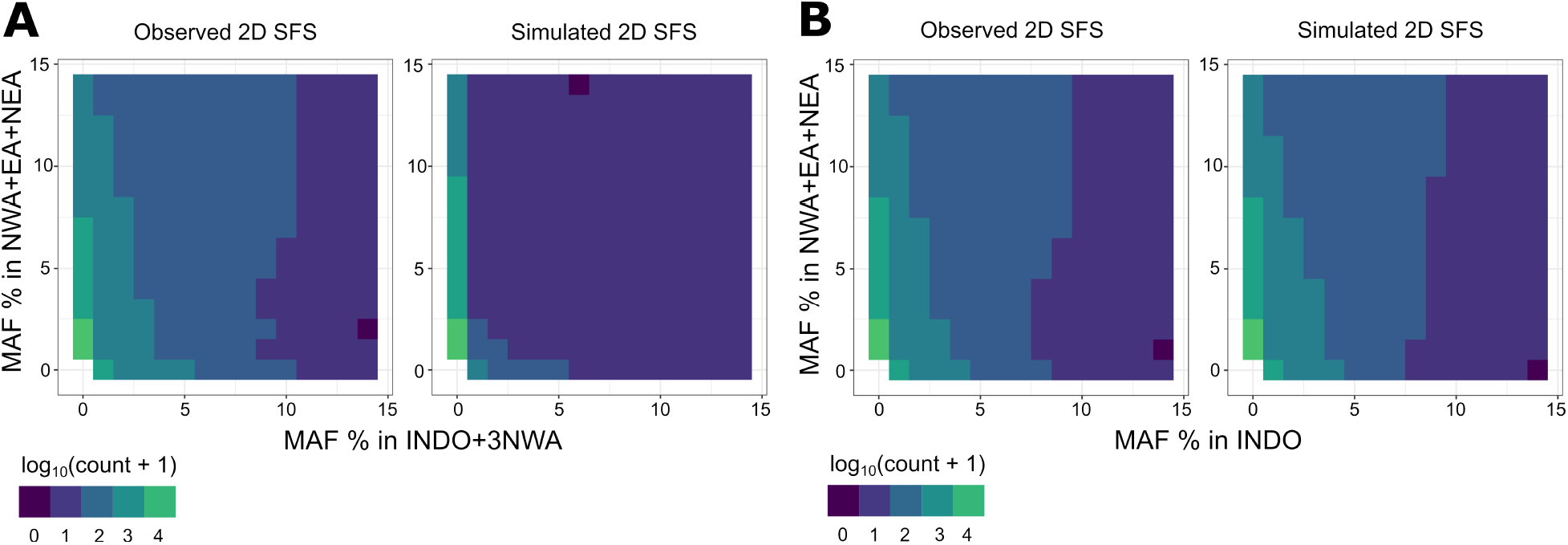
Comparison of the observed and simulated joint 2D site frequency spectrum (SFS) under the best-fit demographic model for Australian and Indonesian *Pteropus alecto*. The SFS was computed using A) all samples and B) excluding the samples from North West Australia genetically clustered with the Indonesian population (samples M53754, M54659, and M54660). Log-transformed counts of sites are shown for minor allele frequency (MAF) pairs. Monomorphic sites are omitted. The following population codes were used: INDO+3NWA, Indonesia and three samples from North West Australia; INDO, Indonesia; NWA+EA+NEA, North West Australia, East Australia and North East Australia.

**Figure S19.**
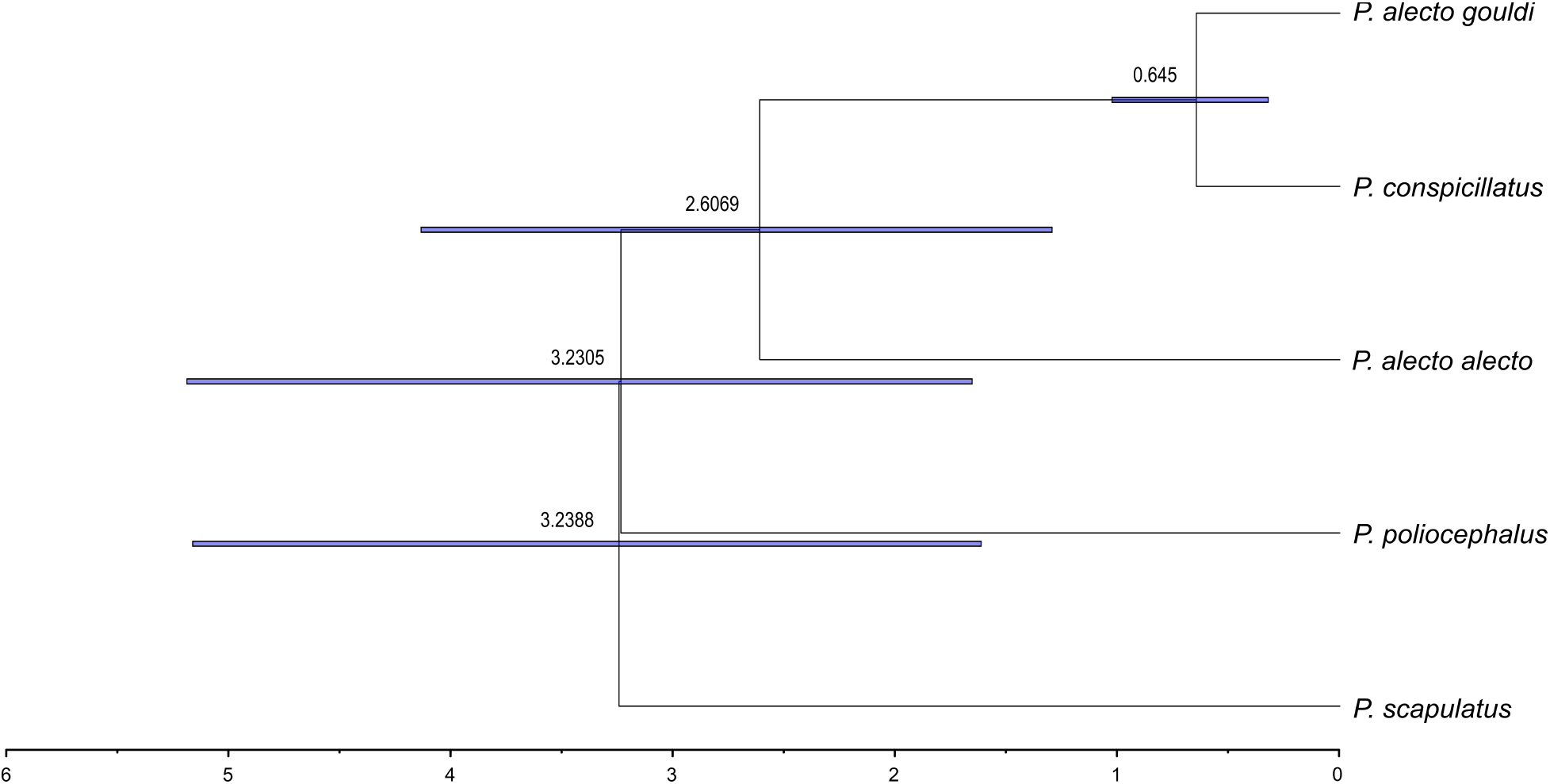
Dated phylogeny of sampled flying-fox species inferred using BEAST and SNAPP from genome-wide SNPs for three samples per branch. Mean divergence estimates per node are ages in millions of years before present. Blue bars show the 95% highest posterior density interval for each node. The subspecies designation *Pteropus alecto gouldi*, used to describe Australian *P. alecto*, is used here to indicate that the three sampled individuals for this branch all originate from Australia.

