## Supplementary text for "Extreme mobility creates contrasting patterns of panmixia, isolation-by-distance and hybridization in four flying-fox species (*Pteropus*)"

### Text S1. DNA extraction and double digest restriction-site-associated DNA sequencing (ddRAD) library preparation

Samples were stored in 80% ethanol at room temperature until processing. Genomic DNA was extracted using the Qiagen DNA Blood and Tissue Kit (Hilden, Germany) with an overnight incubation at 56°C until the tissue was completely lysed. The ddRAD libraries (Peterson et al., 2012) were prepared by digesting a total amount of 100 ng of genomic DNA from each individual in a 50 µL reaction with 20 units each of high-fidelity restriction enzymes, *PstI* and *EcoRI* (New England Biolabs, Ipswich MA, USA), 10× CutSmart buffer and water for 30 min at 37°C, followed by a heat inactivation step for 20 min at 65°C. The digestion products were allowed to slowly cool to room temperature and held at 4°C. The digestion product was ligated to the modified P1 and P2 adapters ensuring that each individual had a unique combination of P1 and P2 barcodes. This way twenty-four P1 and four P2 adapters allowed for multiplexing 96 individuals. The ligation reactions were set up in 70 µL volume using the full volume of digestion product with 4 µL of 50 nM P1 and 12 µL of 50 nM P2 adapters, 2000 units of T4 DNA ligase (New England Biolabs, Ipswich MA, USA), 0.15 mM rATP (Promega, Madison, WI, USA), and water and were incubated for 30 min at 22°C. Ligations were heat inactivated for 20 min at 65°C and allowed to slowly cool to room temperature and held at 4°C. Ten µL of adapter-ligated DNA fragments from all individuals were pooled, with careful mixing after each sample addition. The pooling was replicated in three tubes and the following size-selection steps performed on each tube.

The larger fragments (>300 bp) were extracted with 0.65× volume of Agencourt® AMPure® XP magnetic bead solution (Beckman Coulter, Brea, CA, USA) and the bead discarded. The eluent was transferred to a new tube, cleaned with 0.1× volume of bead solution, washed twice with 200 µL of freshly made 80% ethanol, air-dried and resuspended in 20 µL elution buffer (Qiagen, Hilden, Germany). The three replicates were then pooled and well mixed. Finally, 15 µL of size selected DNA was used as a template in a 60 µL PCR reaction with 30 µL of Phusion High Fidelity 2× Master Mix (New England Biolabs, Ipswich MA, USA), 3 µL of 10 µM P1 (AATGATACGGCGACCACCGAGATCTACACTCTTTCCCTACACGACG) and P2

(CAAGCAGAAGACGGCATACGAGATCGTGATGTGACTGGAGTTCAGACGTGTGC) primers and water. PCR conditions were: 98°C for 30 s, 20 cycles of 98°C for 10 s, 60°C for 30 s, 72°C for 40 s, and the final elongation at 72°C for 5 min. The PCR product was cleaned with a 1.8× bead solution and eluted in 40 µL elution buffer to make the final library.

**Text S2. Sample quality control**

A total of 17 samples failed alignment quality control thresholds requiring a >90% paired alignment rate and >800,000 aligned reads. A single sample was excluded based on genotype missingness >55%. A further 6 samples were flagged for expert review and excluded as contaminants based on discordance of preliminary species assignment with sample metadata including sampling location and museum specimen review. For 22 duplicate samples, the sample with the best alignment metrics was selected for further analysis. All samples and relevant exclusion criteria per sample are listed in Table S1. Two *P. poliocephalus* samples collected outside of the usual range of this species in Ingham (samples 1815076 and 1815091) were confirmed to be classified correctly.

**Text S3. Bayesian divergence time estimation**

To investigate between-species divergence-times and validate parameters inferred with fastsimcoal2, we used BEAST2 2.6.7 (Bouckaert et al., 2019) with the plugin SNAPP to infer divergence-times based on SNPs (Stange et al., 2018). We selected three samples without inferred admixture per species retaining a total of 10,394 variable sites. Ascertainment bias correction to adjust parameters based only on variable sites was used. A lognormal prior (real mean=3.54, sd=0.3, offset=0) on the crown group containing *P. poliocephalus*, *P. alecto* and *P. conspicillatus* was set to calibrate the analysis based on a previously inferred age of 3.54 mya (Almeida et al., 2014). The moderately high standard deviation was selected to account for the uncertainty inherent in secondary calibrations and to ensure overlap with other slightly deviating estimates for the group (Tsang et al., 2019). The previously inferred maximum likelihood tree was used as a fixed topology to constrain the SNAPP analysis. Inference was run for over 40 million MCMC generations with a 10% burn-in and convergence was confirmed using visual inspection in Tracer 1.7.2 with a criterion of effective sample size (ESS) >200 for each parameter.
